# An arbitrary-spectrum spatial visual stimulator for vision research

**DOI:** 10.1101/649566

**Authors:** Katrin Franke, André Maia Chagas, Zhijian Zhao, Maxime J.Y. Zimmermann, Yongrong Qiu, Klaudia Szatko, Tom Baden, Thomas Euler

## Abstract

Visual neuroscientists require accurate control of visual stimulation. However, few stimulator solutions simultaneously offer high spatio-temporal resolution and free control over the spectra of the light sources, because they rely on off-the-shelf technology developed for human trichromatic vision. Importantly, consumer displays fail to drive UV-shifted short wavelength-sensitive photoreceptors, which strongly contribute to visual behaviour in many animals, including mice, zebrafish and fruit flies. Moreover, many non-mammalian species feature more than three spectral photoreceptor types. Here, we present a flexible, spatial visual stimulator with up to 6 arbitrary spectrum chromatic channels. It combines a standard digital light processing engine with open source hard- and software that can be easily adapted to the experimentalist’s needs. We demonstrate the capability of this general visual stimulator experimentally in the *in vitro* mouse retinal whole-mount and the *in vivo* zebrafish. Hereby, we intend starting a community effort of sharing and developing a common stimulator design.

## Introduction

### Challenges in visual stimulation

From psychophysics to single-cell physiology, neuroscientists fundamentally rely on accurate stimulus control. At first glance, generating visual stimuli appears to be much easier than, for example, olfactory stimuli, because computer screens and video projectors are omnipresent, suggesting a range of cost-effective choices for the vision researcher. However, commercially available display devices target human consumers and, thus, are designed for the primate visual system. These devices provide superb spatial resolution, approximately cover the colour space relevant for trichromatic human vision (reviewed in Surridge et al., 2003) and support refresh rates that consider the human flicker-fusion frequency (e.g. Hecht and Verrijp, 1933). Moreover, as the emphasis is on improving the subjective viewing experience, commercial display devices typically lack or even purposefully distort properties that are important when used as visual stimulator for research.

While the spatial resolution provided by even basic commercial displays is typically in excess of what most model species can resolve, limitations may exist with respect to timing (i.e. refresh rate) and, in particular, colour space. For example, many insects including *Drosophila* have flicker fusion frequencies in excess of 100 Hz (Miall, 1978) and use five or more main visual opsins (Feuda et al., 2016). For most vertebrate model species (e.g. mice and zebrafish), standard refresh rates of ~60 Hz suffice for the majority of stimulus requirements, however, the limited colour space poses a serious issue: The light sources (i.e. light-emitting diodes, LEDs) are selected based on the spectrum spanned by the human cone photoreceptor opsins (Dartnall et al., 1983; Nathans et al., 1986) and spectrally arranged to cover the human trichromatic colour space. Hence, these devices fail to generate adequate colours for species with different spectral photoreceptor sensitivities and typically three-channel devices impose further limitations for species with more than three spectral types of (cone) photoreceptor (reviewed in Baden and Osorio, 2018).

Since some of the aforementioned constraints are “hard-wired” in display devices for the consumer market, it is often impractical if not impossible to modify such devices. Specialised solutions aimed to overcome some of the above constraints are commercially available, for instance, as special calibrated LCD monitors for human psychophysics (e.g. Display++, Cambridge Research Systems, Rochester, UK). However, these solutions are expensive, optimised for primates and often closed source, which makes it difficult for the user to modify them. As a result, vision researchers either invest large amounts of time and/or money aiming to overcome these constraints or are forced to settle on a custom suboptimal solution that addresses the needs of a particular experimental situation. This, in turn, may critically limit the stimulus space that can be routinely explored, and yields substantial problems in reproducibility and interpretation when comparing physiological data between laboratories. Comparability and reproducibility are of particular interest in the backdrop of recent developments in increasingly efficient data acquisition technologies. For example, being able to simultaneously record from 100s of neurons using multielectrode arrays (e.g. Jun et al., 2017) or two-photon functional imaging (e.g. Ahrens et al., 2013; Stringer et al., 2018) means that experimental limitations are rapidly shifting the “bottleneck” away from the recording side towards the visual stimulation side.

### Visual stimuli for current animal models

Choosing the adequate animal model for a specific research question may, on one hand, greatly facilitate the experimental design and the interpretation of the results. On the other hand, when trying to transfer such results to other species, it is critical to keep in mind that each species is adapted to different environments and employs different strategies to survive and procreate (reviewed in Baden and Osorio, 2018). In vision research, classical studies often used monkeys and cats as model organisms, which with respect to visual stimuli, e.g. in terms of spatial resolution and spectral sensitivity range, have similar requirements as humans. Today, frequently used animal models -- such as *Drosophila*, zebrafish or rodents -- feature adaptations of their visual systems “outside the specifications” for human vision: For instances, all of the aforementioned species possess UV-sensitive photoreceptors, zebrafish have tetrachromatic vision, and both zebrafish and *Drosophila* display higher flicker fusion frequencies than most mammals (reviewed in Marshall and Arikawa, 2014; Boström et al., 2016). Still, many studies in these species use visual stimulation devices produced and optimized for humans. At best, this will sub-optimally drive the animal model′s visual system, potentially resulting in wrong interpretations of the data.

The visual stimulator we describe in this article can be easily adapted to different animal models. We demonstrate the adaptability of our solution using two exemplary animal species: mice and zebrafish (larvae). Mice currently represent a frequently used model for the mammalian visual system and serve as an example for UV-sensitive vision, while zebrafish are a representative for a well-studied non-mammalian vertebrate species with tetrachromatic vision. Since species-specific chromatic requirements are often more difficult to meet than, for instance, sufficient spatial resolution, our focus here is on adequate chromatic stimulation.

To achieve adequate chromatic stimulation, the spectral composition of the light sources in the stimulator need to cover the spectral sensitivity of the respective model organism. In the ideal case, there should be (*i*) as many LED peaks as the number of spectrally separable photoreceptor types and (*ii*) these should be distributed across the spectral sensitivity range of the species. In general, the spectral sensitivity of an animal is determined by the palette of light-sensitive proteins expressed in their photoreceptors. Vertebrate photoreceptors are divided into rod photoreceptors (rods) and cone photoreceptors (cones). Rods are usually more light-sensitive than cones and, hence, serve vision at dim illumination levels, whereas cones are active at brighter light levels and support colour vision. Depending on the peak sensitivity and the genetic similarity of their opsins, cones are grouped into short (sws, “S”), medium (mws, “M”) and long wavelength-sensitive (lws, “L”) types, with the sws cones further subdivided into sws1 (near-ultraviolet to blue range, <430 nm) and sws2 (blue range, >430 nm) (reviewed in Ebrey and Koutalos, 2001; Yokoyama, 2000). The rod:cone ratio of a species is related to the environmental light levels during their activity periodes. For instance, while the central fovea of the macaque monkey retina lacks rods altogether, the rod:cone ratio in its periphery is approx. 30:1 (Wikler and Rakic, 1990) and therefore similar to that in mice (Jeon et al., 1998). In adult zebrafish, the rod:cone ratio is approx. 2:1 (Hollbach et al., 2015).

Many vertebrates feature a single type of rod (for exceptions, see Baden and Osorio, 2018) but up to 5 spectral types of cone, which is why cones are more relevant for chromatically adequate visual stimulation. Old-world primates including humans, for example, possess three spectral types of cones (S, M and L). Hence, these primates feature trichromatic daylight vision (reviewed in Jacobs, 2008). In contrast, mice are dichromatic like the majority of mammals; they only have two cone types (S and M; Fig. 1a-c). Unlike most mammals, however, the spectral sensitivities of the mouse are shifted towards shorter wavelengths, resulting in a UV-sensitive S-opsin (Jacobs et al., 1991). While one cone type usually expresses only one opsin type, some mammalian species, such as mice or guinea pigs, show opsin co-expression: In mice, for instance, M-cones co-express S-opsin with increasing levels towards the ventral retina (Fig. 1b) (Applebury et al., 2000; Baden et al., 2013; Röhlich et al., 1994). As a “more typical” example for non-mammalian vertebrates, the cone-dominated retina of zebrafish contains four cone types, resulting in tetrachromatic vision (Fig. 1d): In addition to S- and M-cones they have also UV- and L-cones (Chinen et al., 2003). In adult zebrafish, all cone types are organized in a highly regular array, with alternating rows of UV-/S- and M-/L-cones (Fig. 1e,f) (Li et al., 2012). In zebrafish larvae, however, the cone arrangement shows distinct anisotropic distributions for different cone types matched to image statistics present in natural scenes (Fig. 1g,h) (Zimmermann et al., 2018).

**Figure 1.**
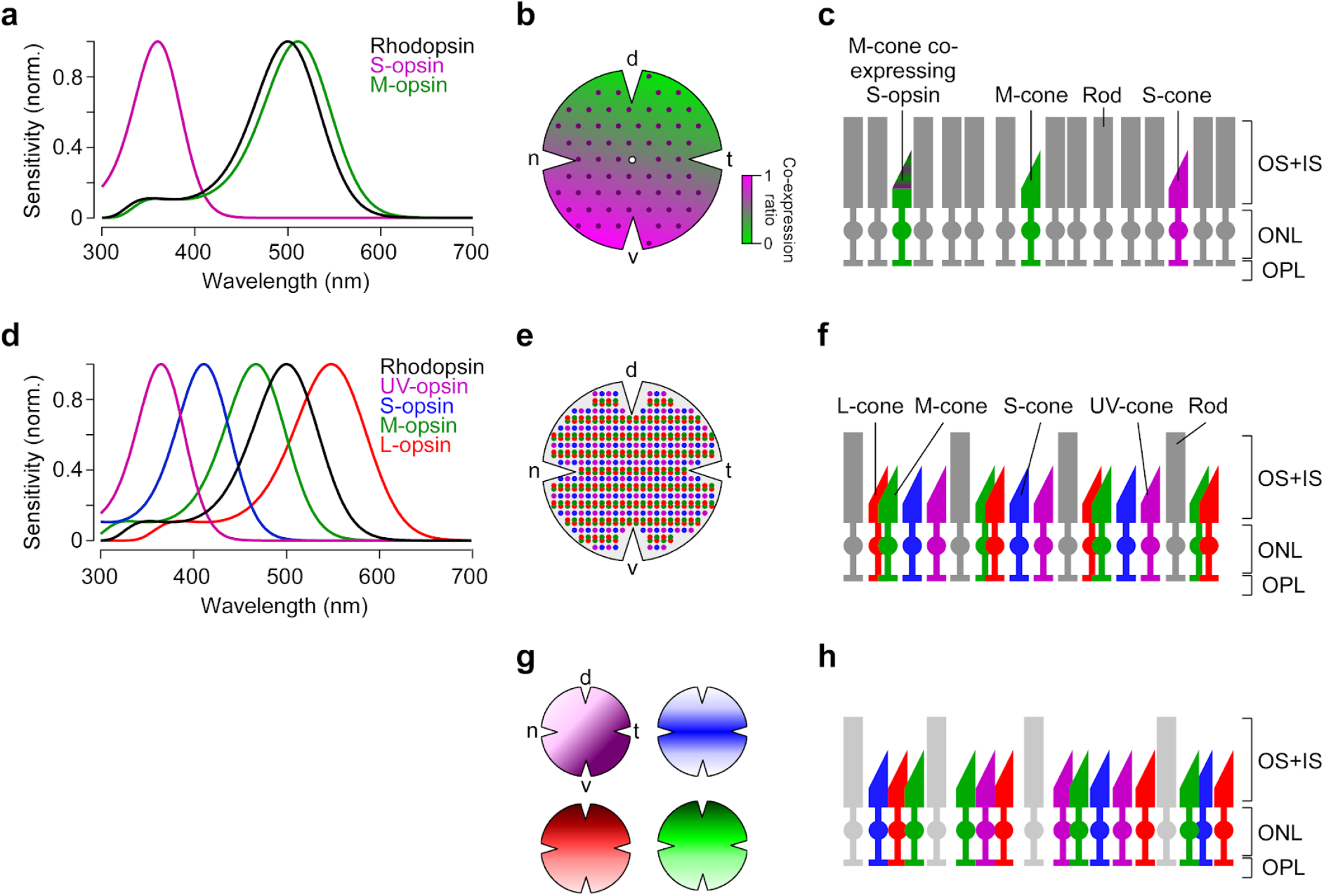
Photoreceptor types and distribution in mouse and zebrafish retina. **a**, Peak-normalized sensitivity profiles of mouse S- (magenta) and M-opsin (green) as well as Rhodopsin (black; profiles were estimated following Stockman and Sharpe, 2000). **b**, Schematic drawing of the distribution of cone photoreceptor (cone) types in the mouse; rod photoreceptors (rods) are homogeneously distributed (Jeon et al., 1998) (not shown here). Purple dots represent “true” S-cones exclusively expressing S-opsin (Haverkamp et al., 2005); ratio of co-expression of S-opsin in M-cones (Applebury et al., 2000; Baden et al., 2013) is colour-coded from green to magenta (d, dorsal; t, temporal; v, ventral; n, nasal). **c**, Illustration of mouse cone and rod arrangement (vertical view; OS+IS, outer and inner segments; ONL, outer nuclear layer; OPL, outer plexiform layer). **d**, Peak-normalized sensitivity profiles of zebrafish UV- (magenta), S- (blue), M- (green) and L-opsin (red) as well as Rhodopsin (black). **e**, Schematic illustration of the regular cone arrangement in adult zebrafish. Coloured dots represent UV-, S-, M- and L-cones. **f**, Like (c) but for adult zebrafish retina. **g**, Schematic drawing illustrating the distribution of cone types in zebrafish larvae (Zimmermann et al., 2018). Colours as in (d). **h**, Like (c,f) for zebrafish larvae. Lighter color of rods indicate that they are not functional at this age (7-9 dpf; Branchek and Bremiller, 1984; Morris and Fadool, 2005).

Taken together, the diversity of spectral sensitivities present in common animal models used in visual neuroscience as well as their differences to the human visual system necessitates a species-specific stimulator design. Here, we present a highly flexible, relatively low-cost visual stimulation system that combines digital light processing (DLP) technology with easily customisable mechanics and electronics, as well as intuitive control software written in Python. We provide a detailed description of the stimulator design and discuss its limitations as well as possible modifications and extensions; all relevant documents are available online (for links, see Table 1). Finally, we demonstrate the use of our stimulator in two exemplary applications; as a dichromatic version for *in vitro* two-photon (2P) recordings in whole-mounted mouse retina and as a tetrachromatic version for *in vivo* 2P imaging in zebrafish larvae.

**Table 1.**
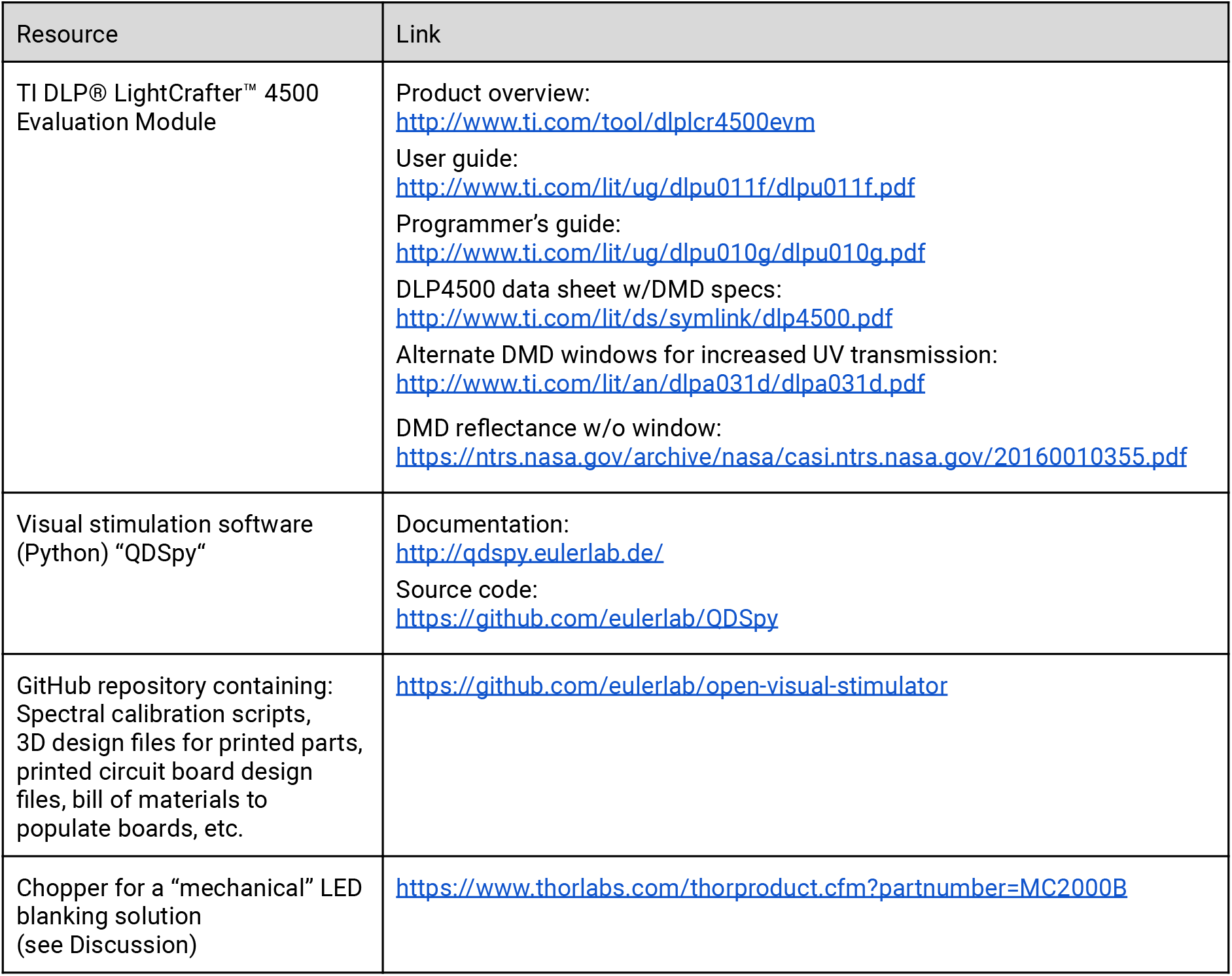
Links to online material.

## Results

### Stimulator design

As the “light engine” of our stimulator, we use the DLP® LightCrafter™ 4500 (here, referred to as “LCr”) developed by Texas Instruments (Dallas, TX, USA). This and similar DLP-based light engines are broadly used in consumer products, such as beamers. The LCr is a bare-metal version for developers and offers several advantages over consumer devices: (*i*) its control protocol is well documented (for links, see Table 1), allowing to program the device via an USB connection on-the-fly; (*ii*) its flexibility in terms of light sources; lightcrafters with customized LEDs and a version with a light guide port are available; (*iii*) its small footprint facilitates incorporating the LCr into existing setups. While the stimulators are built around the LCr, we attempted to use a minimum of commercial parts. Except for the specialised optical elements (i.e. dichroic filters, beam splitters, mirrors), most parts can be replaced by 3D printed designs to increase flexibility and to lower the total costs. For example, instead of commercial rail systems, such as LINOS microbank (Qioptiq, Göttingen, Germany), alternative 3D-printed parts can be used (Delmans and Haseloff, 2018). All electronics and the visual stimulation software are Open Source (Table 1).

For the two-channel (dichromatic) mouse stimulator (Fig. 2a-c), we used a LCr (Fiber-E4500MKIITM, EKB Technologies Ltd., Israel) that was coupled by a light guide to an external illumination unit (Fig. 2a right, c). In this unit, a long-pass dichroic mirror combines the light from two band pass-filtered LEDs (with λ_*peak*_ = 387, 576 nm) and feeds it into a light guide using a collimator (for parts list, see Table 2). This arrangement facilitates the exchange of the LEDs and allows to mount the illumination unit outside the microscope cabinet. One disadvantage with this current LCr model is, however, that -- in our experience -- it passes only a fraction of the light entering the light guide port (see Discussion). The LCr is positioned next to the microscope’s stage and projects the stimulus via a condenser from below into the recording chamber, where it is focussed on the photoreceptor layer of the isolated mouse retina (Fig. 2a left). In the type of two-photon (2P) microscope used (MOM, Sutter Instruments, Novato, CA, USA; Methods), the scan head including the objective lens -- as well as the substage assembly with the condenser -- moves relative to the static recording chamber. Hence, to allow the stimulus to “follow” the objective lens-condenser axis, the LCr is mounted on a pair of low-friction linear slides, with the LCr mechanically coupled to the substage assembly (Fig. 2b). To allow for stimulus centering, a combination of an x-y and a z-stage, both manually adjustable with micrometer screws, is fitted between slides and LCr.

**Table 2.**
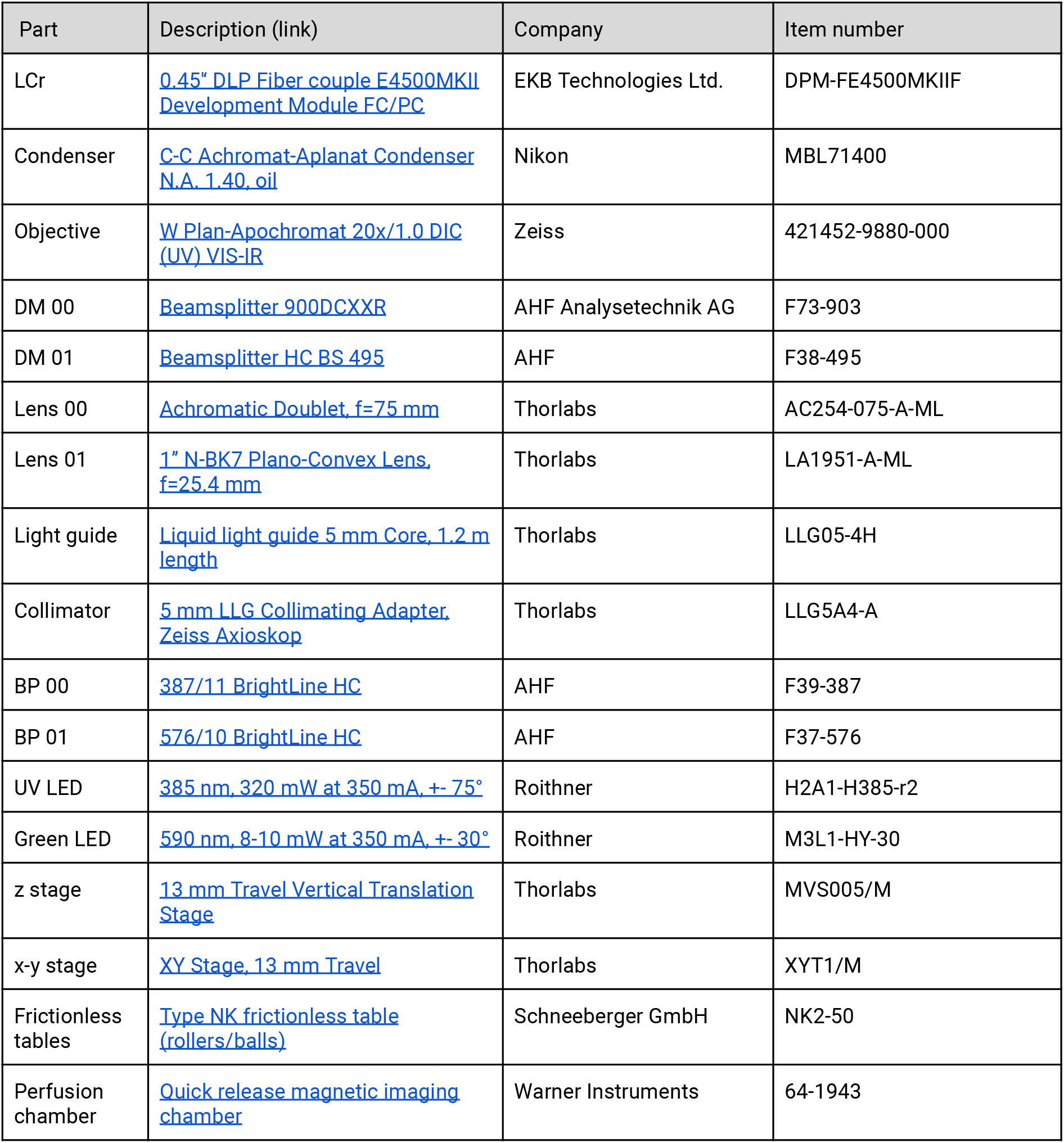
Parts list of the mouse visual stimulator (*cf*. Fig. 2a-c).

**Figure 2.**
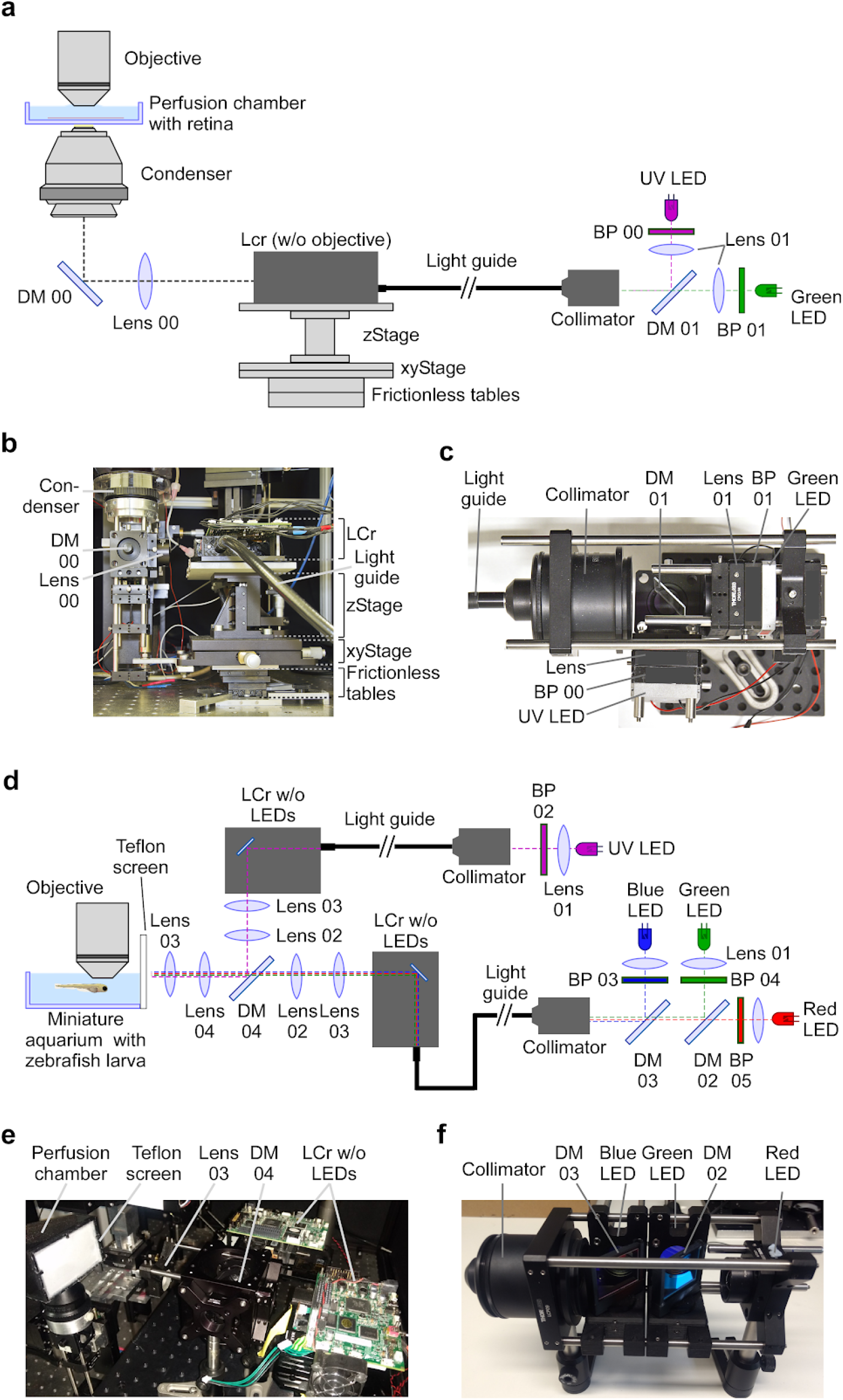
Visual stimulator design. **a**, Schematic drawing of the dichromatic stimulator for *in vitro* recordings of mouse retinal explants. The stimulator is coupled into the two-photon (2P) microscope from below the recording chamber with the retinal tissue (through-the-condenser; for alternative light paths (through-the-objective), see Supplemental Fig. S1). DM, dichroic mirror; BP, band-pass filter; LCr, lightcrafter; LED, light-emitting diode. For specifications of the components, see Table 2. **b,** LCr unit and substage portion of the 2P microscope in side-view. **c,** External LED illumination unit in top-view. **d,** Schematic drawing of the tetrachromatic stimulator for *in-vivo* recordings in zebrafish larvae. The optical pathways of two LCrs are combined and the stimulus is projected onto a UV-transmissive teflon screen at one side of the miniature aquarium. For specifications of the components, see Table 3. **e,** Side-view of tetrachromatic stimulation setup. **f,** “RGB” external LED illumination unit of tetrachromatic stimulation setup. Band-pass (BP) filters 03, 04 and 05 as well as lenses 01 are not indicated due to space constraints. For design of printed parts, see Github repository (Table 1).

**Table 3.**
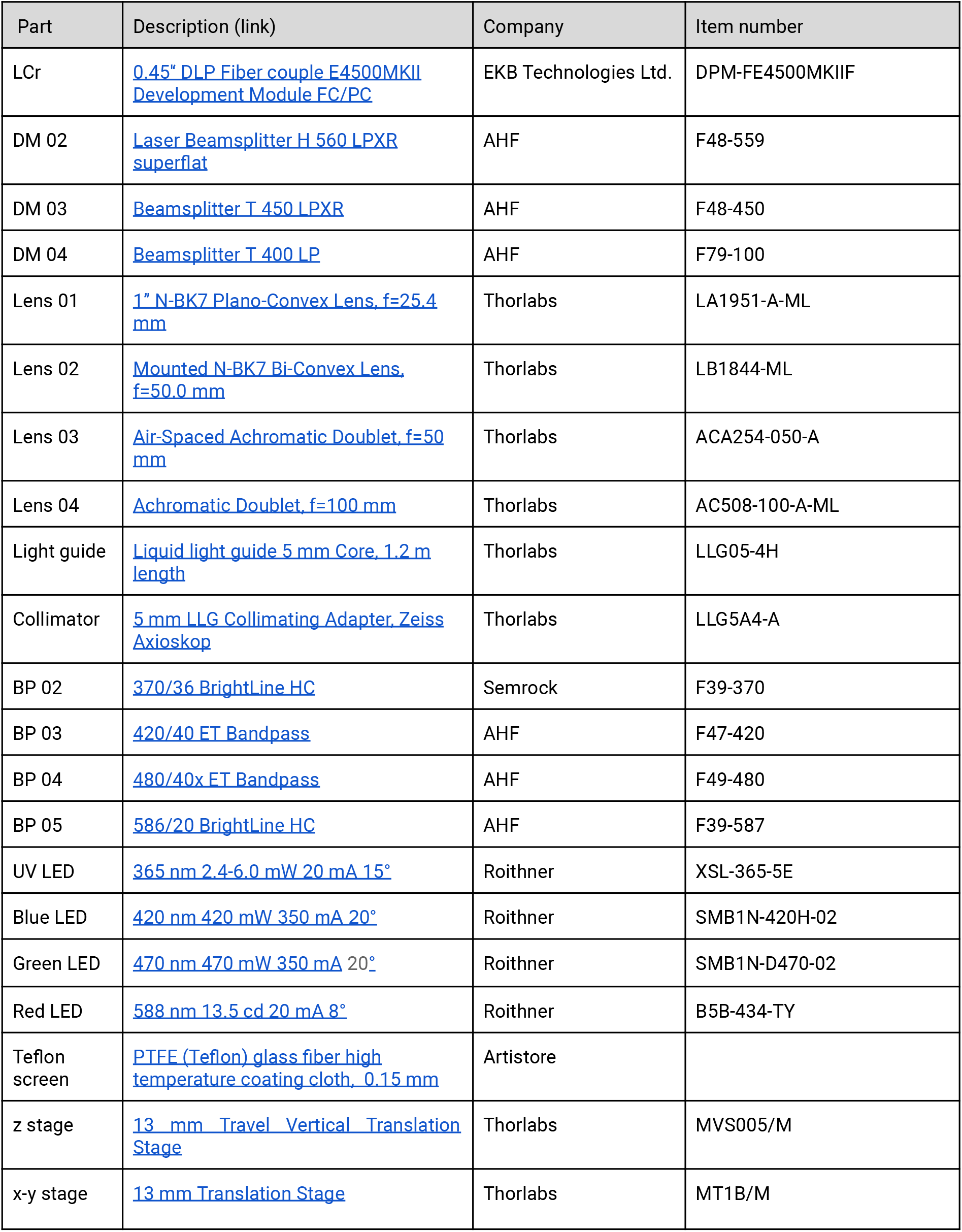
Parts list of the zebrafish visual stimulator (*cf*. Fig. 2d,e). Parts listed for building the external LED unit are not indicated in Fig. 2.

In addition to this “through-the-condenser” (TTC) configuration for the visual stimulation, we also used the “through-the-objective” (TTO) configuration described earlier (Supplemental Fig. S1; Euler et al., 2009). Here, the stimulus is optically coupled into the laser pathway and therefore does not require mechanical coupling of microscope head and visual stimulator. In addition, only scattered light of the visual stimulus will reach the photodetectors above the objective, reducing artifacts caused by stimulus light entering the photomultipliers (PMTs). However, the disadvantage of the TTO configuration is that the stimulation area is limited by the field-of-view of the objective (approx. 700 μm in diameter for our 20x objective) and, therefore, large-scale retinal networks that may be critical for naturalistic stimulation are likely not well activated.

For the **four-channel (tetrachromatic) zebrafish stimulator** variant, we optically coupled two light guide LCrs (Fig. 2d,e; for parts list, see Table 3). They used a similar external illumination unit as the mouse stimulator, but with different LED/filter combinations (λ_*peak*_ = 586, 480, 420, and 370 nm). The beams of the two LCrs are collimated and combined using a long-pass dichroic mirror and projected onto a flat teflon screen that covers one side of a miniature water-filled aquarium (see Table 1), in which the zebrafish larva is mounted on a microscope slide under the objective lens of a MOM-type 2P microscope (Fig. 2e). Each LCrs is mounted on an independent 3-axis manipulator to facilitate alignment of the two images. Then, small 0.5-cm circular stimuli are projected (one from each stimulator) and the LCrs positions are adjusted using the manipulators until the stimuli are completely overlapping.

### Separating light stimulation and fluorescence detection

A difficulty when combining visual stimulation with fluorescence imaging is that the spectral photoreceptor sensitivities and the emission spectra of the fluorescent probes tend to greatly overlap. Hence, to avoid imaging artifacts, stimulator light has to be prevented from reaching the photodetectors (e.g. photomultipliers; PMTs) of the microscope, while ensuring that each of the spectral photoreceptor types is stimulated efficiently and as much of the fluorescence signal as possible is captured. To address this issue, light stimulation and fluorescence detection have to be separated spectrally and/or temporally (Euler et al., 2009). Since dichroic filters are not perfect and photodetectors are extremely light sensitive, we used both approaches in combination.

**Spectral separation** is implemented by designing dichroic mirrors with multiple transmission bands (Euler et al., 2019, 2009), e.g. to transmit one narrow band of stimulation light for each spectral photoreceptor type while reflecting the excitation laser (> 800 nm) and the fluorescence signals (detection bands; Supplemental Fig. S2).

**Temporal separation** means that the LEDs of the visual stimulator are turned off while collecting the fluorescence signal. In a “standard” rectangular x-y image scan, the retrace period (when the scanners move to the beginning of next scan line) can be used for turning the LEDs on to display the stimulus. We found a retrace period of 20% of a scan line (for 1 to 2 ms scan lines) a good compromise between maximising data collection time, avoiding mechanical artifacts from the retracing galvo scanners, and still having sufficient bright stimuli (Euler et al., 2019). Such LED-on periods can also be embedded into more “mechanical scanner friendly” scan patterns that require little or no retrace time, such as spiral scans (Rogerson et al., 2018). In either case, for this to work the microscope’s software has to signal these “retrace” periods. Our microscope software (ScanM, see Material) generates a “laser blanking signal” (=low during retrace), which allows turning down the excitation laser’s intensity via a Pockels’ cell during retrace to reduce the laser exposure of the tissue (Euler et al., 2019, 2009). Hence, a straightforward way to implement temporal separation between fluorescence detection and light stimulation is to invert this blanking signal and use it to turn on the LEDs during retrace (Supplemental Figure S3a,b).

The LCr contains a single DMD chip and generates coloured images by sequentially presenting the R/G/B contents (as 3×8 bitplanes) of each frame while turning on the respective LED driver (*cf*. LCr User Guide; for link, see Table 1). The intensity of a, say, green pixel is defined by the temporal pattern the corresponding DMD-mirror is flicked between the “on” and the “off” position while the green LED is constantly on. In addition, the LCr allows to set the maximal intensity of each LED via pulse-width modulation (PWM). In the two-channel mouse stimulator, we power the LEDs in the external illumination unit (Fig. 2a right) using the LCr’s onboard LED drivers and therefore, these LEDs are driven as built-in ones would be -- except that we interrupt power to the LEDs in sync with the inverted laser blanking signal using a simple custom circuit (Supplemental Fig. S4a-c). To switch the necessary currents with sufficient speed, this circuit uses 3 solid state relays connected in parallel. For the four-channel zebrafish stimulator, we devised a more general solution that uses only the LCr’s digital control signals for each LED (LED enable, LED PWM; for details, see Supplemental Fig. S4d-g). This solution has the advantage that it can use an arbitrary combination of LED, LED driver and power supply. For all solutions, printed circuit board (PCB) designs and building instructions are provided (see link to repository in Table 1).

One potential issue of the described solution for temporal separation is that the frame (refresh) rate of the LCr (typically 60 Hz) and the laser blanking / LED-on signal (500 to 1,000 Hz) are not synchronised and therefore may cause slow aliasing-related fluctuations in stimulus brightness. In practice, however, we detected only small brightness modulations(Supplemental Fig. S3c).

### Visual stimulation software

Our visual stimulation software (QDSpy) is completely written in Python 3 and relies on OpenGL for stimulus rendering. Each stimulus is a Python script that calls the “QDSpy library” to define stimulus objects, set colors, send trigger signals, display scenes etc. Stimulus objects range from simple shapes with basic shader support to videos (for a complete description of the software, follow link in Table 1). The QDSpy GUI facilitates spatial stimulus alignment, LCr control and stimulus presentation. When a stimulus script is executed for the first time, it generates a “compiled” version of the stimulus, which is then used by the software when presenting the stimulus. The main advantage of this procedure is a reliable timing as potential delays in the execution of the Python script are minimized; its main disadvantage is that user interaction during stimulus presentation is (currently) not possible.

For stimulus presentation, QDSpy relies on the frame sync of the graphics card/driver for stimulus display. By measuring the time required to generate the next frame, the software can detect dropped frames and warn the user of timing inconsistencies. Such frame drops, including all other relevant events (e.g. which stimulus was started when, was it aborted etc.) as well as user comments are automatically logged into a file. Currently, we do not reprogram the LCr firmware and, instead, run it in “video mode”, where the LCr behaves like a standard HDMI-compatible display (60 Hz, 912×1140 pixels). To account for any gamma correction performed by the LCr firmware when in video mode and/or by non-linearities of the LEDs /LED drivers, we measured each LED’s intensity curve separately to generate a lookup table (LUT) that is then used in QDSpy to linearize the colour channels (Methods).

The stimulation software allows generating digital synchronisation markers to align presented stimuli with recorded data. In addition to digital I/O cards (e.g. PCI-DIO24, Measurement Computing, Bietigheim-Bissingen, Germany), QDSpy supports Arduino boards (https://www.arduino.cc/) as digital output device. While the software attempts generating the synchronisation marker at the same time as when presenting the stimulus frame that contains the marker, a temporal offset between these two events in the 10s of millisecond range cannot be avoided. We found this offset to be constant for a given stimulation system, but dependent on the specific combination of PC hardware, digital I/O device, and graphic cards. Therefore, the offset must be measured (e.g. by comparing synchronization marker signal and LCr output measured by a fast photodiode) and considered in the data analysis.

For up to 3 chromatic channels (e.g. the mouse stimulator, *cf*. Fig. 2a-c), stimuli are presented in full-screen mode on the LCr, with the other screen displaying the GUI. When more chromatic channels are needed, as for the zebrafish stimulator, two LCrs are combined (see above; *cf.* Fig. 2d,e). QDSpy then opens a large window that covers both LCr “screens” and provides each LCr with “its” chromatic version of the stimulus (“screen overlay mode”). To this end, the software accepts color definitions with up to 6 chromatic values and assigns them to the 6 available LEDs (3 per LCr). For example, the first LCr of the zebrafish stimulator provides the red, green and blue channels, whereas the second LCr adds the UV channel (Fig. 2d). Here, QDSpy presents the stimulus’ RGB-components on the half of the overlay window assigned to the first LCr and the stimulus’ UV-component on the half of the overlay window assigned to the second LCr. The remaining LED channels are available for a different purpose, such as, for example, separate optogenetic stimulation.

### LED selection and spectral calibration

Adequate chromatic stimulation requires adjusting the stimulator to the spectral sensitivities of the model organism. Ideally, one would choose LEDs that allow maximally separating the different opsins (Fig. 3). In practice, however, these choices are limited by the substantial overlap of opsin sensitivity spectra (Fig. 3a,c) and by technical constraints: For instance, commercially available projectors, including the LCr, barely transmit UV light (<385 nm), likely due to UV non-transmissive parts in the optical pathway and/or the reflectance properties of the DMD (Discussion). In addition, when imaging light-evoked neural activity, fluorescence signal detection and visual stimulation often compete for similar spectral bands, and need to be separated to avoid stimulus-related artefacts (Supplemental Fig. S2; discussed in Euler et al., 2019, 2009). Because LED spectra can be quite broad, we combine each LED with an appropriate band-pass filter to facilitate arranging stimulation and detection bands.

**Figure 3.**
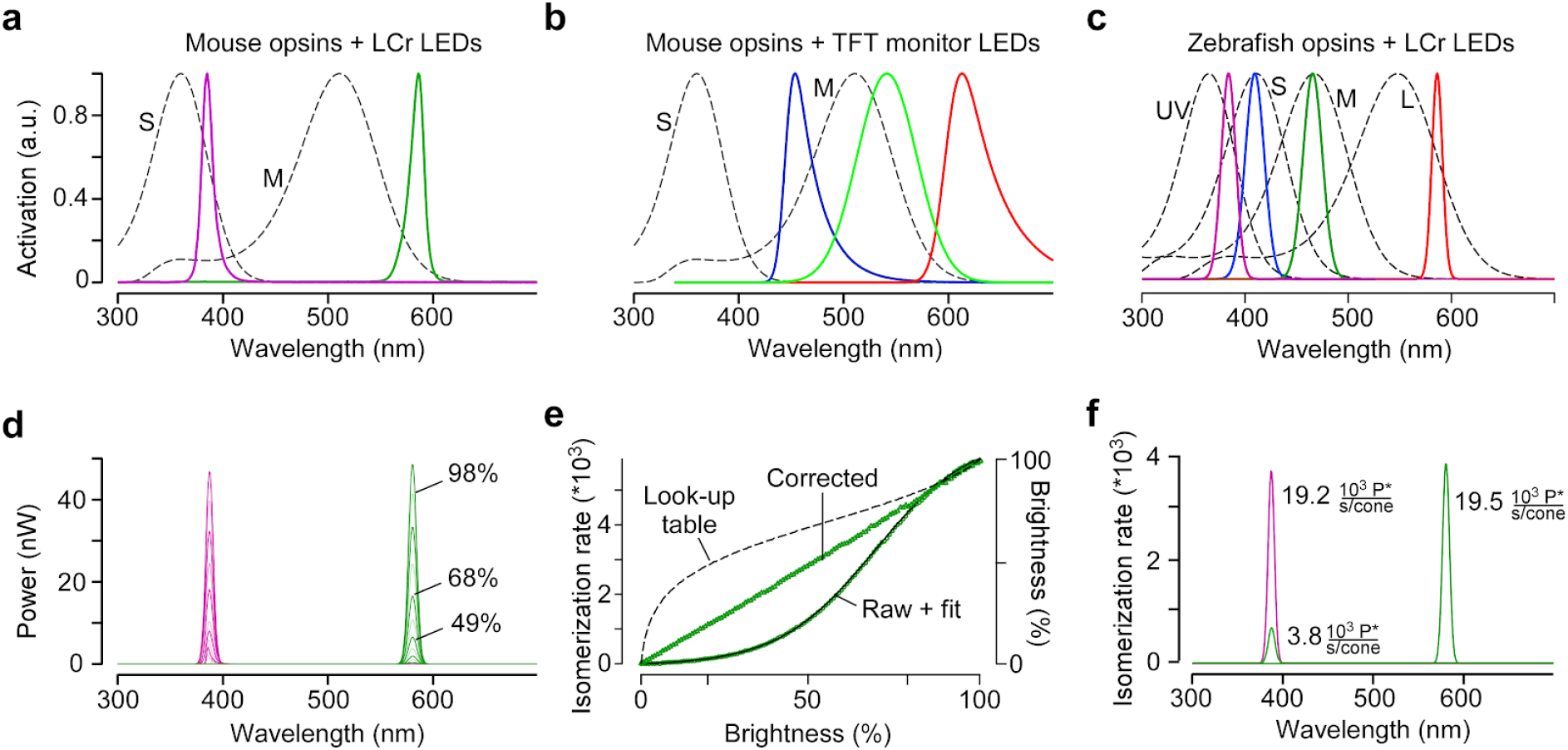
Calibration of the mouse stimulator. **a,** Sensitivity profiles of mouse S- and M-opsin (dotted black lines) and spectra of UV (magenta) and green LED/filter combinations used in the mouse stimulator. **b,** Sensitivity profiles of mouse S- and M-opsin (dotted black line) and spectra of blue, green and red LED present in a standard TFT monitor. **c,** Sensitivity profiles of zebrafish opsins (dotted black lines) and spectra of UV, blue, green and red LEDs used in the zebrafish stimulator. **d,** Spectra (in nW) of UV and green LED obtained from measurements using increasing brightness levels; shown are spectra for 0, 9, 19, 29, 39, 49, 59, 68, 78, 88, and 98% brightness. **e,** Non-linearized intensity curve (“raw”) with sigmoidal fit (black), estimated gamma correction curve (black dotted line; “Look-up table”) and linearized intensity curve (“corrected”) for green LED. **f,** Photoisomerization rates for maximal brightness of UV (19.2 and 3.8 photoisomerizations/s for S- and M-opsin, respectively) and green LED (0 and 19.5 photoisomerizations/s for S- and M-opsin, respectively). Note that the UV LED also activates M-opsin due to its increased sensitivity in the short wavelength range (β-band, Discussion).

As a consequence, the peak emissions of the selected LED/filter combinations usually do not match the opsins’ sensitivity peaks. For our dichromatic mouse stimulator, we chose LED/filter combinations peaking for UV and green at approx. 385 and 576 nm, respectively (Fig. 3a), which after calibration (Fig. 3d-f; Methods), are expected to differentially activate mouse M- and S-opsin (Fig. 3f). Notably, because of its spectral shift towards shorter wavelengths (Jacobs et al., 1991), conventional TFT monitors routinely used in *in-vivo* studies fail to activate mouse S-opsin (Fig. 3b) and therefore are not able to provide adequate visual stimuli for the mouse visual system (Discussion). For the tetrachromatic zebrafish stimulator, we used LED/filter combinations with peak emissions at approx. 586, 480, 420, and 370 nm (Fig. 3c).

To estimate the theoretically achievable chromatic separation of mouse cones with our stimulators, we measured the spectra of each LED/filter combination at different intensities (Fig. 3d) and converted these data into cone photoisomerisation rates (Nikonov et al., 2006). To account for non-linearities in stimulator intensities, we apply gamma correction at the stimulus presentation software level (Fig. 3e). Details on these procedures and example calculations for mice and zebrafish are provided in the Methods and in supplemental iPython notebooks, respectively (Table 1).

### Spatial resolution

To measure the spatial resolution of our **mouse stimulator**, we used the “through-the-objective” (TTO) configuration (Supplemental Fig. S1; Euler et al., 2009) and projected UV and green checkerboards of varying checker sizes (from 2 to 100 μm; Methods) onto a camera chip positioned at the level of the recording chamber (Fig. 4a). We found that contrast remained relatively constant for checker sizes down to 4 μm before it rapidly declined (Fig. 4b,c). Similarly, transitions between bright and dark checkers started to blur for checker sizes below 10 μm (Fig. 4d,e). For these measurements, we used a 5x objective (MPlan 5X/0.1, Olympus) to project the stimuli, ensuring that the spatial resolution of the camera (OVD5647 chip: 1.4 μm pixel pitch) was not the limiting factor. Hence, a 5 × 5 μm checker stimulus appeared as a 20 × 20 μm square on the camera chip, where it covered approx. (14.3)^2^ pixels. However, for the scaling factor we use for our recordings (1.9 × 0.9 μm/pixel), a 5 × 5 μm checker consists only of 9.5 × 4.5 LCr pixels (DMD4500, chip area: 6,161 × 9,855 μm with 1,140 × 912 pixels). Thus, the drop in spatial resolution observed for checkers ≤5 μm is likely related to the resolution of the DMD.

**Figure 4.**
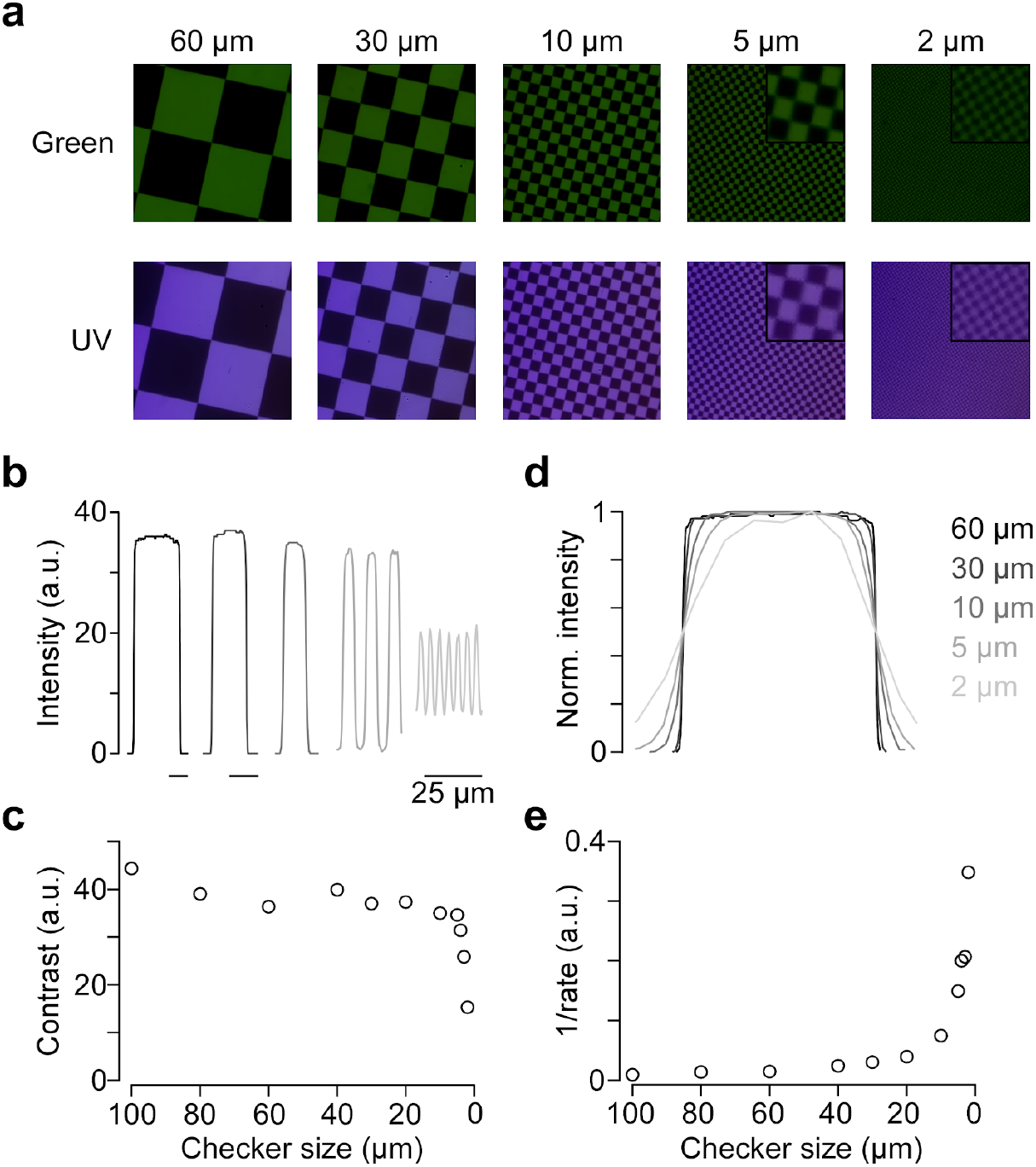
Spatial calibration of the mouse stimulator. **a,** Images of checkerboard stimuli with varying checker sizes for illumination with green (top) and UV (bottom) LED, recorded by placing the sensor chip of a raspberry pi camera at the level of the recording chamber. Focus was adjusted for UV and green LED separately. Insets for 5 and 2 μm show zoomed in regions of the image. **b,** Intensity profiles for five different checker sizes of green LED. **c,** Contrasts (*I*_*Max*_ − *I*_*Min*_) for checkerboards of varying checker sizes. **d,** Peak-normalized intensity profiles of different checker sizes, scaled to the same half-maximum width. **e,** 1/rate estimated from sigmoidal fits of curves in (d) for varying checker sizes.

For the spatial resolution measurements, the UV and green images were each focussed on the camera chip and, therefore, the results do not reflect any effects of chromatic aberration on image quality. To estimate chromatic aberration for our TTO configuration, we next measured the offset between the focal planes of the chromatic channels. Here, we used the standard 20x objective that we also employ for functional recordings. We found that the difference in focal plane between UV and green of approx. 24 μm has little effect on the overall image quality (Supplemental Fig. S5) - at least for checker sizes we routinely use for receptive field mapping of retinal neurons (e.g. Baden et al., 2016; *cf*. also Discussion).

### Visual stimulation in the explanted mouse retina

To confirm that our stimulator design can be used for adequate chromatic stimulation of the mouse retina, we directly recorded from cone photoreceptor axon terminals in retinal slices using 2P Ca^2+^ imaging (e.g. Kemmler et al., 2014). To this end, we used the transgenic mouse line HR2.1:TN-XL, where the ratiometric Ca^2+^ sensor TN-XL is exclusively expressed in cones (Fig. 5a) (Wei et al., 2012). To quantify the chromatic preference of recorded cones, we calculated spectral contrast (*SC*) based on the response strength to a 1 Hz sine-wave full-field stimulus of green and UV LED (Methods). The *SC* values correspond to Michelson contrast, ranging from −1 to 1 for the cell responding solely to UV and green, respectively.

**Figure 5.**
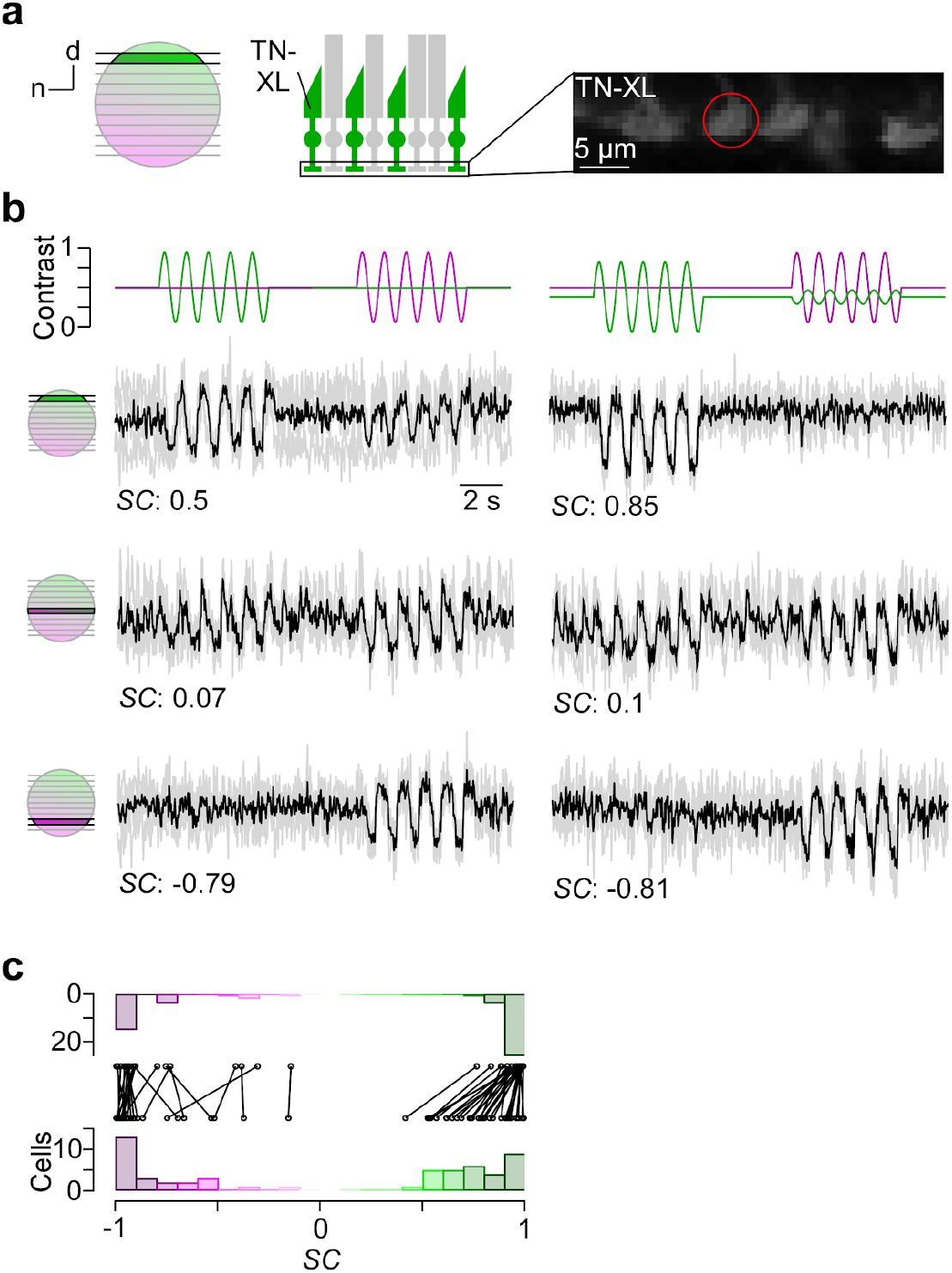
Cone-isolating stimulation of mouse cones. **a,** Dorsal recording field in the outer plexiform layer (OPL; right) shows labeling of cone axon terminals with Ca^2+^ biosensor TN-XL in the HR2.1:TN-XL mouse line (Wei et al., 2012). Schematic on the left illustrates retinal location of recorded slice. **b,** Ca^2+^ traces (mean traces in black, n=3 trials in grey) of cone axon terminals located in dorsal (top; cone axon terminal from (a)), medial (middle) and ventral (bottom) retina in response to 1 Hz sine modulation of green and UV LED, with spectral contrast (*SC*) indicated below. Color substitution protocol (right) estimated from calibration data (Methods). **c,** Distribution and comparison of *SC* for sine modulation stimulus with (top) and without (bottom) silent substitution protocol (n=55 cells, n=12 scan fields, n=1 mouse; p<0.01 for dorsal cells, n=30; p>0.05 for ventral cells, n=25; Wilcoxon signed-rank test).

In line with the opsin distribution described in mice (Applebury et al., 2000; Baden et al., 2013), cones located in the ventral retina responded more strongly or even exclusively to UV (Fig. 5b, bottom row), whereas central cones showed a strong response to both green and UV due to the more balanced co-expression of S- and M-opsin (Fig. 5b, center row). In contrast, dorsal cones exhibited a green-dominated response (Fig. 5b, top row). Due to the cross-activation of M-opsin by the UV LED (see above), most dorsal cones showed an additional small response to UV.

We also tested a stimulus that used silent substitution (Fig. 5b, right column; Methods) (Estévez and Spekreijse, 1982). With this stimulus, we systematically found reduced UV responses in dorsal cones, resulting in a significant shift in *SC* towards more positive values (Fig. 5c, right column; for statistics, see legend). In contrast, ventral cone responses were not altered by silent substitution.

These data demonstrate that our stimulator design enables obtaining cone-isolating responses in mouse retina. Notably, the chromatic separation observed in the recordings nicely matches our predictions of cross-activation (see above and Methods).

### Tetrachromatic stimulation in in-vivo zebrafish larvae

We recorded *in vivo* from bipolar cell (BC) axon terminals in zebrafish larvae using 2P Ca^2+^ imaging (Fig. 6a). The transgenic line we used expressed SyGCaMP6f exclusively in BC axon terminals (Dreosti et al., 2009). In these experiments, we presented full-field (90 × 120 degrees visual angle) steps or sine wave modulation of red, green, blue and UV light to the teflon screen in front of the immobilized animal (*cf*. Fig. 2d,e). This revealed spectrally differential tuning of distinct BC terminals (Fig. 6b,c), in line with a previous report (Zimmermann et al., 2018). For example, terminal 1 responded with a Ca^2+^ increase to a decrease in red light as well as to an increase in blue or UV light, yielding a “red^Off^/blue^On^,UV^On^” response behaviour. In contrast, terminal 4 did not respond to red or green, but differentially responded to blue and UV (“blue^Off^/UV^On^”). Further differences were visible in the temporal profile of the BC responses. For example, terminal 3 responded more transiently to red and blue, but in a sustained fashion to UV. Similar to cone responses in the *in vitro* mouse retina, spectrally differential tuning of zebrafish BC terminals was also observed for a sine wave stimulus (Fig. 6c). Taken together, tetrachromatic stimulation elicited clear differential responses across different wavelengths, thus highlighting that the stimulator’s spectral isolation between the four LED channels was sufficient to drive the zebrafish’s cone system differentially.

**Figure 6.**
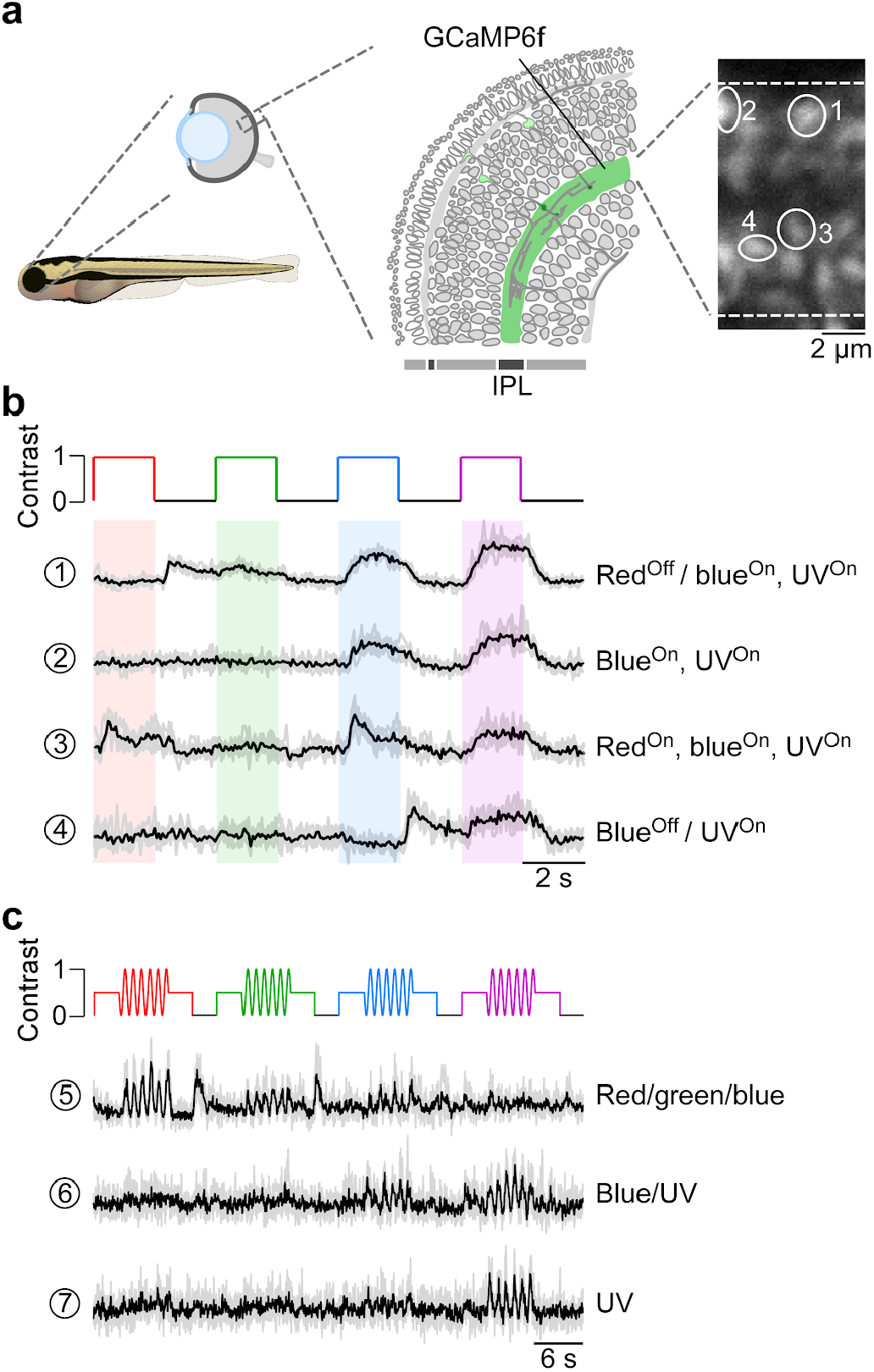
Chromatic responses in bipolar cells of *in vivo* zebrafish larvae. **a,** Drawing illustrating the expression of the genetically encoded Ca^2+^ biosensor SyGCaMP6f in bipolar cell terminals (left) of *tg(1.8ctbp2:SyGCaMP6f)* zebrafish larvae and scan field of inner plexiform layer (IPL; right), with exemplary ROIs marked by white circles. **b,** Mean Ca^2+^ traces (black; n=6 trials in grey) in response to red, green, blue and UV full-field flashes (90 × 120 degrees visual angle, presented to the fish’s right side). **c,** Mean Ca^2+^ traces (black; n=4 trials in grey) in response to full-field sine modulation (at 1 Hz) of red, green, blue and UV LED.

## Discussion

In this paper, we present a flexible, relatively low-cost stimulator solution for visual neuroscience and demonstrate its use for dichromatic stimulation in the *in vitro* mouse retina and tetrachromatic stimulation in the *in vivo* larval zebrafish. The core of the stimulator is a LCr with a custom LED complement or, for more flexibility, a light guide port that connects to an external LED array. We also provide detailed calibration protocols (as iPython notebooks) to estimate (cross-)activation in a species’ complement of photoreceptor types, which facilitates planning of the LED/filter combinations required for selective chromatic stimulation. To drive the LEDs, we designed simple electronic circuits that make use of the LCr LED control signals and allow integrating an LED-on signal (“blanking signal”) for synchronisation with data acquisition, which is critical, for example, for fluorescence imaging in the *in vitro* retina (Euler et al., 2009). By combining two LCrs, up to 6 LED channels are supported by our visual stimulation software (QDSpy). In addition, we describe three exemplary projection methods that allow tuning the system towards high spatial resolution (“through-the-objective”) or a large field-of-stimulation (“through-the-condenser”) for *in vitro* experiments, or presentation on a teflon screen for *in vivo* studies. All material (electronics, optical design, software, parts lists etc.) are publically available and open source.

### The need for “correct” spectral stimulation

The spectral sensitivity markedly varies across common model organisms used in visual neuroscience (*cf*. Introduction). As a result, in most cases visual stimulation devices optimized for the human visual system do not allow “correct” spectral stimulation, in the sense that the different photoreceptor types are not differentially activated by the stimulator LEDs. Instead, “correct” spectral stimulation requires that the visual stimulator is well-adjusted to the specific spectral sensitivities of the model organism.

For example, while human S-opsin is blue-sensitive (reviewed in Jacobs, 2008), the S-opsin of mice shows its highest sensitivity in the UV range (Fig. 1b) (Jacobs et al., 1991). As standard TFT monitors optimized for humans and routinely used in mouse *in vivo* studies do not emit in the UV range, they fail to activate mouse S-opsin (*cf.* Fig. 3b). If then, due to space constraints, the stimulation monitors are positioned in the UV-sensitive upper visual field of the mouse (*cf.* Fig. 1b), such a stimulator will mainly activate the rod pathway. As a result, the presentation of “truely” mouse-relevant natural stimuli is hampered, if not impossible. In recent years, however, several studies used customized projectors that allow UV stimulation for investigating chromatic processing in, for example, dLGN (Denman et al., 2017) or V1 (Tan et al., 2015). Here, similar to the arrangement in our zebrafish stimulator (*cf.* Fig. 2d), the image is either back-projected onto a UV-transmissive teflon screen (Tan et al., 2015) or projected onto a visual dome coated with UV-reflective paint (Denman et al., 2017). Both solutions are compatible with the mouse stimulator described above.

Even when the stimulator is adjusted to the spectral sensitivity of the model organism, each stimulator LED typically activates more than one photoreceptor type due to overlapping sensitivity profiles of the different opsins (*cf.* Fig. 1). In particular, the long sensitivity tail of opsins for shorter wavelengths (“β-band”) contributes to cross-activation of photoreceptors by the stimulator LEDs. For example, the sensitivity of mouse M-opsin to our UV LED results in a cross-activation of ~19.5% (*cf.* Fig. 3f). Such “imperfect” spectral separation of cone types is sufficient to investigate many questions concerning chromatic processing in the visual system -- especially as there rarely is photoreceptor type-isolating stimulation in natural scenes (Chiao et al., 2000). If needed, photoreceptor cross-activation can be ameliorated by using a silent substitution protocol (Estévez and Spekreijse, 1982). Here, one type of photoreceptor is selectively stimulated by presenting a steady excitation to all other photoreceptor types using a counteracting stimulus (*cf*. Fig. 5b). This allows, for instance, to investigate the role of individual photoreceptor types in visual processing.

### Stimulation with UV light

Sensitivity to UV light is widespread across animal species (reviewed in Cronin and Bok, 2016). Sometimes UV sensitivity may represent a specialised sensory channel; e.g. many insects and potentially some fish use UV-sensitive photoreceptors to detect polarisation patterns in the sky for orientation (Parkyn and Hawryshyn, 1993; Seliger et al., 1994; Wehner, 2001). In most cases, however, UV sensitivity seems to be simply incorporated into colour vision, extending the spectral range accessible to the species. Here, UV sensitivity can play an important role in invertebrate and vertebrate behaviour, including navigation and orientation, predator and prey detection, as well as communication (reviewed in Cronin and Bok, 2016).

For instance, mice possess a UV-sensitive S-opsin, which is co-expressed by M-cones predominantly in the ventral retina (Applebury et al., 2000; Baden et al., 2013). As the ventral retina observes the sky, it was proposed that the ventral UV sensitivity promotes detection of predatory birds, which appear as dark silhouettes against the sky (Calderone and Jacobs, 1995). As UV light dominates the (clear) sky due to increased Rayleigh scattering of short wavelengths, contrasts tend to be higher in the UV channel (discussed in (Applebury et al., 2000; Baden et al., 2013; Cronin and Bok, 2016)). In support of this, recordings from mouse cones suggest that, ventral S/M-cones prefer dark contrasts, whereas dorsal M-cones encode bright and dark contrasts symmetrically (Baden et al., 2013).

Zebrafish larvae express the UV-sensitive sws2 opsin in their UV-cones. UV-vision in zebrafish is likely used for several tasks, including prey detection, predator and obstacle avoidance as well as colour vision (Yoshimatsu et al., 2018; Zimmermann et al., 2018). Like in mice, the distribution of UV-cones is non-uniform across the retinal surface. UV-cone density is highest in the temporo-ventral retina which surveys the upper-frontal part of visual space. This UV-specific *area centralis* is likely a specialisation for prey capture: Larval zebrafish feed on small, water-borne microorganisms such as paramecia, which are largely translucent at long wavelengths of light but readily scatter UV (Johnsen and Widder, 2001; Novales Flamarique, 2013). Next, unlike for most terrestrial animals, predators may appear in any part of visual space in the aquatic environment, and zebrafish invest in UV-dark detection of predator-silhouettes throughout visual space (Losey et al., 1999; Zimmermann et al., 2018). Finally, UV-sensitivity is integrated into retinal circuits for color vision to impact tetrachromatic vision, as originally demonstrated for goldfish (Neumeyer, 1992).

Taken together, to approach natural conditions when probing a UV-sensitive species’ visual system, UV stimulation must be included. Nonetheless, there are some pitfalls specifically linked to UV light stimulation. One major issue is that, in our experience, the standard LCr barely transmits wavelengths <385 nm. As the reflectance of the micromirrors (aluminum) drops only <300 nm and the glass window covering the DMD transmits ≥90% of the light down to 350 nm (see links in Table 1), one limiting factor appears to arise from the LCr optics. Therefore, if shorter wavelengths are required, replacing the internal optics of the projector may be necessary (e.g. Tan et al., 2015). If the different stimulation wavelengths are spread across a large range (e.g. Δλ = 191 and 200 nm for zebrafish and mouse stimulator, respectively; *cf.* Fig. 3a,c), chromatic aberration may become an issue, causing an offset between the focal planes of the different colour channels (*cf*. Supplemental Fig. S5). For our TTO stimulator configuration, we found a focus difference between UV and green in the order of a few tens of micrometers. For a checker size that is commonly used for receptive field mapping of retinal neurons (e.g. 40 μm; Baden et al., 2016; Franke et al., 2017), we observed only a slight image blurring due to chromatic aberration, that likely has a negligible effect on our experiments. If chromatic aberration becomes an issue, viable approaches may be to increase the depth-of-field (e.g. by decreasing the aperture size with a diaphragm in the stimulation pathway) and/or use appropriate achromatic lenses.

### Potential issues and technical improvements

In this section, we discuss potential issues that may arise when adapting our stimulator design to other experimental situations, as well as possible technical improvements.

If too much stimulation light enters the PMTs, temporal separation of visual stimulation and data acquisition is needed. To address this problem, we presented here an electronic solution that allows the LEDs to be on only during the short retrace period of a scan line. However, if higher LED power and/or shorter LED-on intervals are needed, the design of the “blanking” circuits becomes more demanding. Here, an alternative is to use a mechanical chopper (see Table 1). Briefly, a custom 3D printed chopping blade is attached to the chopper and the system is mounted at an appropriate position in the light path such that the blade is able to block the stimulus during the system’s scanning period. The blanking signal from the microscope software (see Results) is used to synchronize chopper rotation speed. The main advantage of this solution is that it works with any stimulator and without meddling with its electronics. Disadvantages include, however, (*i*) mechanical vibrations and spinning noise, (*ii*) that different scanning modes require different chopping blades, and (*iii*) the additional costs for the chopper.

For increased flexibility with respect to the LED complement of the visual stimulator, we here use an external LED unit coupled into the LCr via a light guide port (*cf*. Fig. 2). One disadvantage with this LCr model is, however, that it passes only a relatively small fraction of the light entering the light guide port. While this is not problematic for small projection areas used in our mouse and zebrafish recordings or for relatively low light intensities, it may become an issue when projecting the stimulus onto a larger area like the inside of a dome (e.g. Denman et al., 2017; Schmidt-Hieber and Häusser, 2013). Here, the LCr model with built-in, high-power LEDs might be a better option (Supplemental Table 1).

If high spatial resolution is not required, an interesting alternative to a projector-based stimulator is one build from arrays of LEDs (e.g. Reiser and Dickinson, 2008; Wang et al., 2019). The main advantage of LED arrays is that they offer a more precise timing control compared to the combination of HDMI display and PC graphics card driven by software running on a desktop PC. Hence, LED arrays may allow refresh rates in the range of several hundreds of Hz (Reiser and Dickinson, 2008). However, apart from their lower spatial resolution, current LED arrays only support a low number of color channels, making them less well suitable for chromatic processing studies. In addition, LED arrays typically require customized control electronics, whereas stimulators based on standard HDMI displays can be driven by the experimenter’s software of choice.

While we run the LCr for simplicity in “video mode”, where it acts as a normal 60-Hz HDMI display, it is also possible to program the LCr’s firmware in “pattern mode”. Here, the user can precisely define how the incoming stream of RGB bitplanes is interpreted and displayed. For example, if a lower bit depth is sufficient, much higher frame rates are achievable. Texas Instruments provides the documentation of the LCr’s programming interface online (for further details, see links in Table 1).

## Material and Methods

Note that the general stimulator design, operation and performance testing is described in the results section. The respective parts for the mouse and the zebrafish stimulator versions are listed in Tables 2 and 3, respectively. Hence, this section focuses on details about the calibration procedures, 2P imaging, animal procedures, and data analysis.

### Intensity calibration and gamma correction

The purpose of the intensity calibration is to ensure that each LED evokes a similarly maximal photoisomerisation rate in its respective spectral cone type, whereas the gamma correction aims at linearising each LED’s intensity curve. All calibration procedures are described in detail in the iPython notebooks included in the open-visual-stimulator GitHub repository (for link, see Table 1).

In case of the **mouse stimulator**, we used a photo-spectrometer (USB2000, 350-1000 nm, Ocean Optics, Ostfildern, Germany) that can be controlled and read-out from the iPython notebooks. It was coupled by an optic fibre and a cosine corrector (FOV 180°, 3.9 mm aperture) to the bottom of the recording chamber of the 2P microscope and positioned approximately in the stimulator’s focal plane. For intensity calibration, we displayed a bright spot (1,000 μm in diameter, max. intensity) of green and UV light to obtain spectra of the respective LEDs. We used a long integration time (1 s) and fitted the average of several reads (n=10 for green; n=50 reads for UV) with a Gaussian to remove shot noise. This yielded reliable measurements also at low LED intensities, which was particularly critical for UV LEDs.

The spectrometer output (*S*_*meas*_) was divided by the integration time (Δ*t*, in s) to obtain counts/s and then converted into electrical power (*P*_*el*_, in nW) using the calibration data (*S*_*Cal*_, in μJ/count) provided by Ocean Optics,

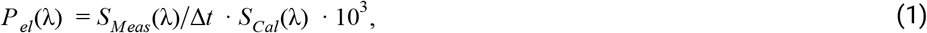

with wavelength λ. To obtain the photoisomerization rate per photoreceptor type, we first converted from electrical power into energy flux (*P*_*eflux*_, in eV/s),

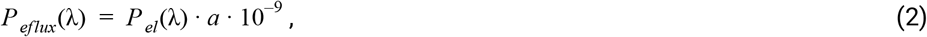

where *a* = 6.242 · 10^18^ eV/J. Next, we calculated the photon flux (*P*_*Phi*_, in photons/s) using the photon energy Q (*P*_*Q*_, in eV),

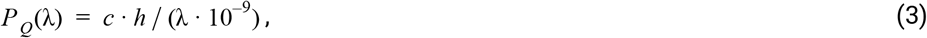

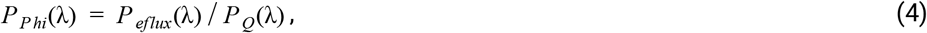

with the speed of light, *c* = 299, 792, 458 m/s, and Planck’s constant, *h* = 4.135667 · 10^−15^ eV⋅s. The photon flux density (*P*_*E*_, [photons/s/μm^2^]) was then computed as

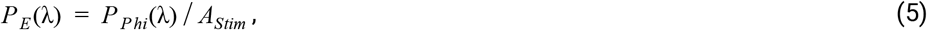

where *A*_*Stim*_ (in μm^2^) corresponds to the light stimulus area. To convert *P*_*E*_ into photoisomerization rate, we next determined the effective activation (*S*_*Act*_) of mouse photoreceptor types by the LEDs as

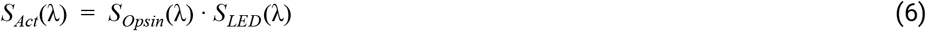

with the peak-normalized spectra of the M- and S-opsins, *S*_*Opsin*_, and the green and UV LEDs, *S*_*LED*_. Sensitivity spectra of mouse opsins were derived from equation 8 in (Stockman and Sharpe, 2000).

For our LEDs (Table 2), the effective mouse M-opsin activation was 14.9% and 10.5% for the green and UV LED, respectively. The mouse S-opsin is only expected to be activated by the UV LED (52.9%) (Fig. 3a,f). Next, we estimated the photon flux (*R*_*Ph*_, [photons/s]) for each photoreceptor as

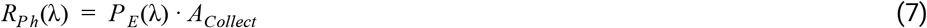

where *A*_*Collect*_ = 0.2 μm^2^ corresponds to the light collection area of cone outer segments (Nikonov et al., 2006). The photoisomerization rate (*R*_*Iso*_, P*/photoreceptor/s) for each combination of LED and photoreceptor type was estimated using

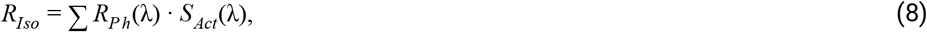

see Nikonov et al. (2006) for details. The intensities of the mouse stimulator LEDs were manually adjusted (Supplemental Fig. S2b-c) to an approx. equal photoisomerisation range from (in P*/cone/s ⋅10^3^) 0.6 and 0.7 (stimulator shows black image) to 19.5 and 19.2 (stimulator shows white image) for M- and S-opsins, respectively (*cf*. Fig. 3f). This corresponds to the low photopic range. The M-opsin sensitivity spectrum displays a “tail” in the short wavelength range (due to the opsin’s β-band, see Fig. 1a and Stockman and Sharpe, 2000), which means that it should be cross-activated by our UV LED. Specifically, while S-opsin should be solely activated by the UV LED (19.2 by UV vs. 0.1 by green; in P*/cone/s ⋅10^3^), we expect M-opsin to be activated by both LEDs (19.5 by green vs. 3.8 by UV). The effect of such cross-activation can be addressed, for instance, by silent substitution (see below).

To account for the non-linearity of the stimulator output using gamma correction, we recorded spectra for each LED for different intensities (1,000 μm spot diameter; pixel values from 0 to 254 in steps of 2) and estimated the photoisomerization rates, as described above. From these data, we computed a lookup table (LUT) that allows the visual stimulus software (QDSpy) to linearize the intensity functions of each LED (*cf*. Fig. 3e; for details, see iPython notebooks; Table 1).

In case of the **zebrafish stimulator**, to determine the LEDs spectra, we used a compact CCD Spectrometer (CCS200/M, Thorlabs, Dachau, Germany) in combination with the Thorlabs Optical Spectrum Analyzers (OSA) software and coupled to a linear fibre patch cable. To determine the electrical power (*P*_*el*_, in nW), we used an optical energy power (PM100D, Thorlabs) in combination with the Thorlabs Optical Power Monitor (OPM) software and coupled to a photodiode power sensor (S130VC, Thorlabs). Both probes were positioned behind the teflon screen (0.15 mm, for details, see Table 3). Following the same procedure, we determined the photoisomerization rate (*R*_*Iso*_, P*/photoreceptor/s) for each combination of LED and photoreceptor types (*cf.* iPython notebooks; Table 1).

### Spatial resolution measurements

To measure the spatial resolution of the mouse stimulator, we removed lens and glass window of a raspberry pi camera chip (OV5647, Eckstein GmbH, Clausthal-Zellerfeld, Germany) and positioned it at the level of the recording chamber. Then, we projected UV and green checkerboards of varying checker sizes (2, 3, 4, 5, 10, 20, 30, 40, 60, 80, and 100 μm) through an objective lense (MPL5XBD (5x), Olympus, Germany) onto the chip of the camera (Fig. 4a). For each checker size and LED, we extracted intensity profiles using ImageJ (Fig. 4b) and estimated the respective contrast as *I*_*Max*_ − *I*_*Min*_ (Fig. 4c). To quantify the steepness of the transition between bright and dark checkers, we peak-normalized the intensity profiles and normalized relative to half-width of the maximum (Fig. 4d). Next, we fitted a sigmoid to the rising phase of the intensity profile

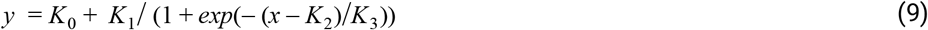

and used 1/*K*_3_ as estimate of the rise time and as a proxy for “sharpness” of the transitions between black and white pixels.

To measure the difference in focal plane of UV and green LED due to chromatic aberration, we projected a 40 and 100 μm checkerboard through a 20x objective (W Plan-Apochromat 20× /1.0 DIC M27, Zeiss, Oberkochen, Germany) onto the raspberry pi camera (see above).

### Photodiode measurements

To measure fast intensity changes of the LEDs of the mouse stimulator, we used a photodiode (Siemens silicon photodiode BPW 21, Reichelt, Sande, Germany; as light-dependent current source in a transimpedance amplifier circuit), which was positioned under a 20x objective (see above). Then, intensity traces in response to green and UV light flashes (10 s) or a full-field chirp stimulus were recorded together with the blanking signal at 250 kHz using pClamp (Molecular Devices, Biberach an der Riss, Germany). To estimate the amount of intensity modulation of the LEDs due to aliasing, we box-smoothed the intensity traces with a box width of 100 ms, which roughly corresponds to the integration time of mouse cone photoreceptors (Umino et al., 2008).

### Silent substitution

For our measurements in mouse cones, we used a silent substitution protocol (Estévez and Spekreijse, 1982) for generating opsin-isolating stimuli to account for the cross-activation of mouse M-opsin by the UV LED. Here, one opsin type is selectively stimulated by presenting a scaled, counterphase version of the stimulus to all other opsin types (*cf*. Fig. 5). Specifically, we first used the ratio of activation (as photoisomerization rate) of M-ospin by UV and green to estimate the amount of cross-activation (*S*_*CrossAct*_). For our recordings, an activation of M-opsin of 19.5 and 3.8 P*/cone/s ⋅10^3^ for green and UV LED resulted in a cross-activation of *S*_*CrossAct*_ = 0.195. Then, *S*_*CrossAct*_ was used to scale the intensity of the counterphase stimulus:

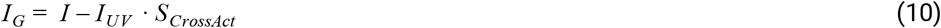

For our recordings in **zebrafish larvae**, we did not use silent substitution. However, we described a possible approach for the zebrafish (or a comparably tetrachromatic species) in our online resources (Table 1).

### Animals and tissue preparation

All animal procedures adhered to the laws governing animal experimentation issued by the German Government (mouse) or all procedures were performed in accordance with the UK Animals (Scientific Procedures) act 1986 and approved by the animal welfare committee of the University of Sussex (zebrafish larvae).

For the **mouse** experiments, we used one 12-week-old HR2.1:TN-XL mouse; this mouse line expresses the ratiometric Ca^2+^ biosensor TN-XL under the cone-specific HR2.1 promoter and allows measuring light-evoked Ca^2+^ responses in cone synaptic terminals (Wei et al., 2012). Animals were housed under a standard 12-h day-night rhythm. Before the recordings, the mouse was dark-adapted for ≥1 h, then anaesthetized with isoflurane (Baxter, Unterschleißheim, Germany) and killed by cervical dislocation. The eyes were removed and hemisected in carboxygenated (95% O_2_, 5% CO_2_) artificial cerebrospinal fluid (ACSF) solution containing (in mM): 125 NaCl, 2.5 KCl, 2 CaCl_2_, 1 MgCl_2_, 1.25 NaH_2_PO_4_, 26 NaHCO_3_, 20 glucose, and 0.5 L-glutamine (pH 7.4). The retina was separated from the eye-cup, cut in half, flattened, and mounted photoreceptor side-up on a nitrocellulose membrane (0.8 mm pore size, Merck Millipore, Darmstadt, Germany). Using a custom-made slicer (Wei et al., 2012; Werblin, 1978), acute vertical slices (200 μm thick) were cut parallel to the naso-temporal axis. Slices attached to filter paper were transferred on individual glass coverslips, fixed using high vacuum grease and kept in a storing chamber at room temperature for later use. For all imaging experiments, individual retinal slices were transferred to the recording chamber of the 2P microscope (see below), where they were continuously perfused with warmed (36°C), carboxygenated extracellular solution.

For the **zebrafish larvae** experiments, we used 7 day post fertilization (*dpf)* larvae of the zebrafish (*Danio rerio)* lines *tg(1.8ctbp2:SyGCaMP6f)*, which expresses the genetically encoded Ca^2+^ indicator GCaMP6f fused with synaptophysin under the RibeyeA promoter and allows measuring light-evoked Ca^2+^ responses in bipolar cell synaptic terminals (Dreosti et al., 2009; Johnston et al., 2019; Rosa et al., 2016; Zimmermann et al., 2018). Animals were grown from 10 hours post fertilization (*hpf)* in 200 μM of 1-phenyl-2-thiourea (Sigma) to prevent melanogenesis (Karlsson et al., 2001). Animals were housed under a standard 14/10-h day-night rhythm and fed 3x a day. Before the recordings, zebrafish larvae were immobilised in 2% low-melting-point agarose (Fischer Scientific, Loughborough, UK; Cat: BP1360-100), placed on a glass coverslip and submersed in fish water. To prevent eye movement during recordings α-bungarotoxin (1 nl of 2 mg/ml; Tocris, Bristol, UK; Cat: 2133) was injected into the ocular muscles behind the eye.

### Two-photon imaging

For all imaging experiments we used MOM-type two-photon (2P) microscopes (designed by W. Denk, MPI, Heidelberg; purchased from Sutter Instruments/Science Products, Hofheim, Germany). For image acquisition, we used custom-made software (ScanM by M. Müller and T.E.) running under IGOR Pro 6.3 for Windows (Wavemetrics, Lake Oswego, OR, USA). The microscopes were equipped each with a mode-locked Ti:Sapphire laser (MaiTai-HP DeepSee, Newport Spectra-Physics, Darmstadt, Germany; or Chameleon Vision-S, Coherent; Ely, UK), two fluorescence detection channels for eCFP (FRET donor; HQ 483/32, AHF, Tübingen, Germany) and citrine (FRET acceptor; HQ 538/50, AHF) or GCaMP6f/GCaMP7b (ET 525/70 or ET 525/50, AHF), and a water immersion objective (W Plan-Apochromat 20× /1.0 DIC M27, Zeiss, Oberkochen, Germany). The excitation laser was tuned to 860 nm and 927 nm for TN-XL (eCFP) in mouse and GCaMP variants in zebrafish, respectively. Time-lapsed image series were recorded with 64 × 16 pixels (at 31.25 Hz) or 128 × 64 (at 15.625 Hz). Detailed descriptions of the setups for mouse (Euler et al., 2019, 2009; Franke et al., 2017) and zebrafish (Zimmermann et al., 2018) have been published elsewhere.

### Data analysis

Data analysis was performed using IGOR Pro (Wavemetrics). Regions of interest (ROIs) of individual synaptic terminals (of mouse cones and zebrafish bipolar cells) were manually placed. Then, Ca^2+^ traces for each ROI were extracted for mouse cones as Δ*R*/*R*, with the ratio *R* = *F*_*A*_/*F*_*D*_ of the FRET acceptor (citrine) and donor (eCFP) fluorescence, and resampled at 500 Hz. For zebrafish bipolar cells, Ca^2+^ traces for each ROI were extracted and detrended by high-pass filtering above ~0.1Hz and followed by z-normalisation based on the time interval 1-6 seconds at the beginning of recordings using custom-written routines under IGOR Pro. A stimulus synchronisation marker that was generated by the visual stimulation software (Results) and embedded in the recordings served to align the Ca^2+^ traces relative to the stimulus with ≤2 ms precision (depending on the scan line duration, see Results and Euler et al., 2019). For this, the timing for each ROI was corrected for sub-frame time-offsets related to the scanning.

#### Response quality index

To measure how well a cell responded to the sine wave stimulus, we computed the signal-to-noise ratio

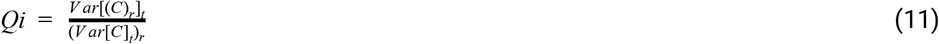

where *C* is the *T* by *R* response matrix (time samples by stimulus repetitions), while ()_*x*_ and *Var*[]_*x*_ denote the mean and variance across the indicated dimension, respectively (Baden et al., 2016; Franke et al., 2017). For further analysis, we used only cells that had a *Qi* > 0.3.

#### Spectral contrast

The mean trace in response to the green and UV sine wave stimulus was used to analyse the spectral sensitivity of the cones. For that, we computed the power spectrum of the trace and used the power (*P*) at the fundamental frequency (1 Hz) as a measure of response strength. Then, the spectral contrast (*SC*) was estimated as

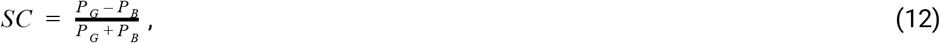

where *P*_*G*_ and *P*_*B*_ correspond to the responses to green and UV, respectively. For statistical comparison of *SC* values with and without silent substitution (see above), we used the Wilcoxon signed-rank test for non-parametric, paired samples.

## Acknowledgment

This study is part of the research program of the Bernstein Center for Computational Neuroscience, Tübingen, and was funded by the German Federal Ministry of Education and Research and the Max Planck Society (BMBF, FKZ: 01GQ1002, and MPG M.FE.A.KYBE0004 to K.F.), the European Research Council (ERC-StG ‘‘NeuroVisEco’’ 677687 to T.B.) the EU’s Horizon 2020 research and innovation programme under the Marie Skłodowska-Curie grant agreement No 674901 (“switchBoard”, to T.B., T.E.), the UKRI (BBSRC, BB/R014817/1 and MRC, MC_PC_15071 to T.B.), the Leverhulme Trust (PLP-2017-005 to T.B.), the Lister Institute for Preventive Medicine (to T.B.), and the Deutsche Forschungsgemeinschaft (DFG, German Research Foundation; Projektnummer 276693517 – SFB 1233 to T.E.).

## Supplemental Information

**Supplemental Fig. S1.**
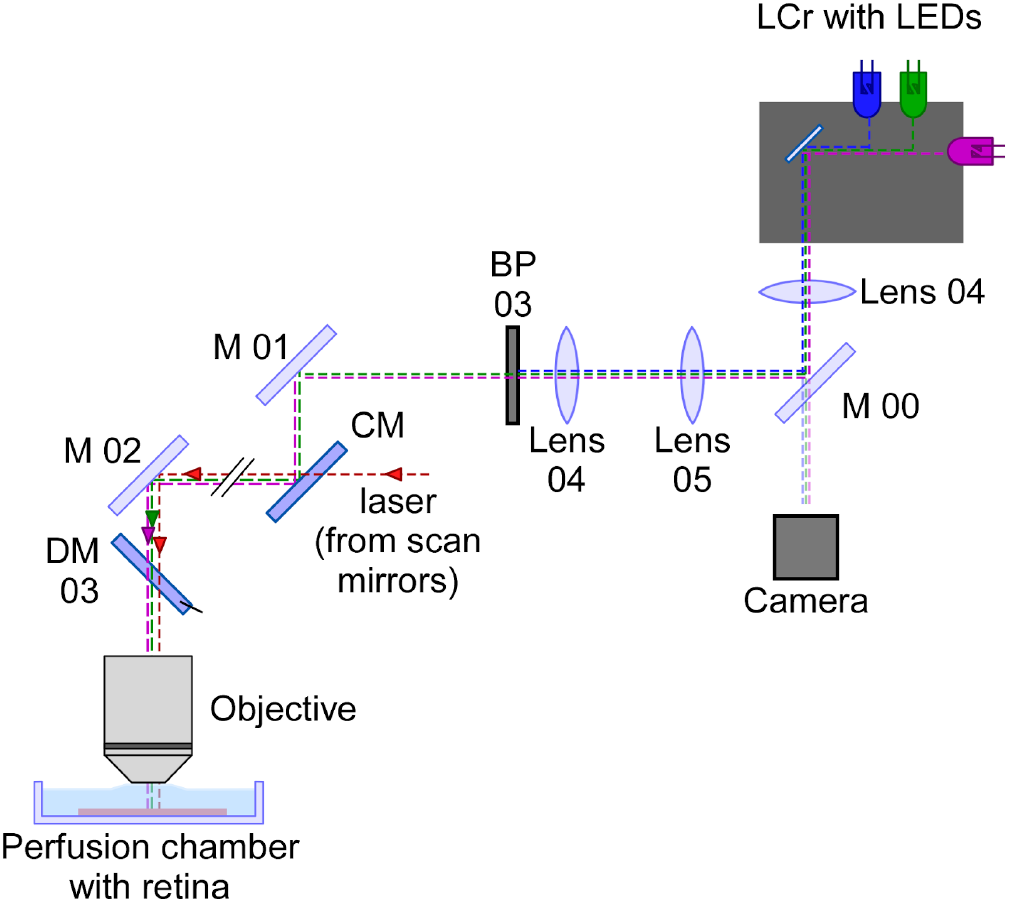
Optical pathway for a through-the-objective (TTO) mouse stimulator. Schematic drawing of a TTO dichromatic stimulator for *in vitro* recordings of mouse retinal explants (cf. Euler et al., 2009). The light from the LCr stimulator with internal UV, blue and green LEDs is filtered by a dual-band filter transmitting UV and green light. Then, the light is coupled into the two-photon microscope using a cold mirror (CM). Using a beam-splitter (M 00), a small fraction of light is projected onto a camera to allow online visualization of the visual stimulus. LCr, lightcrafter; LED, light-emitting diode; M, mirror; CM, cold mirror; DM, dichroic mirror; BP, band-pass filter. For specifications of the components, see Table 2 and Supplemental Table 1.

**Supplemental Fig. S2.**
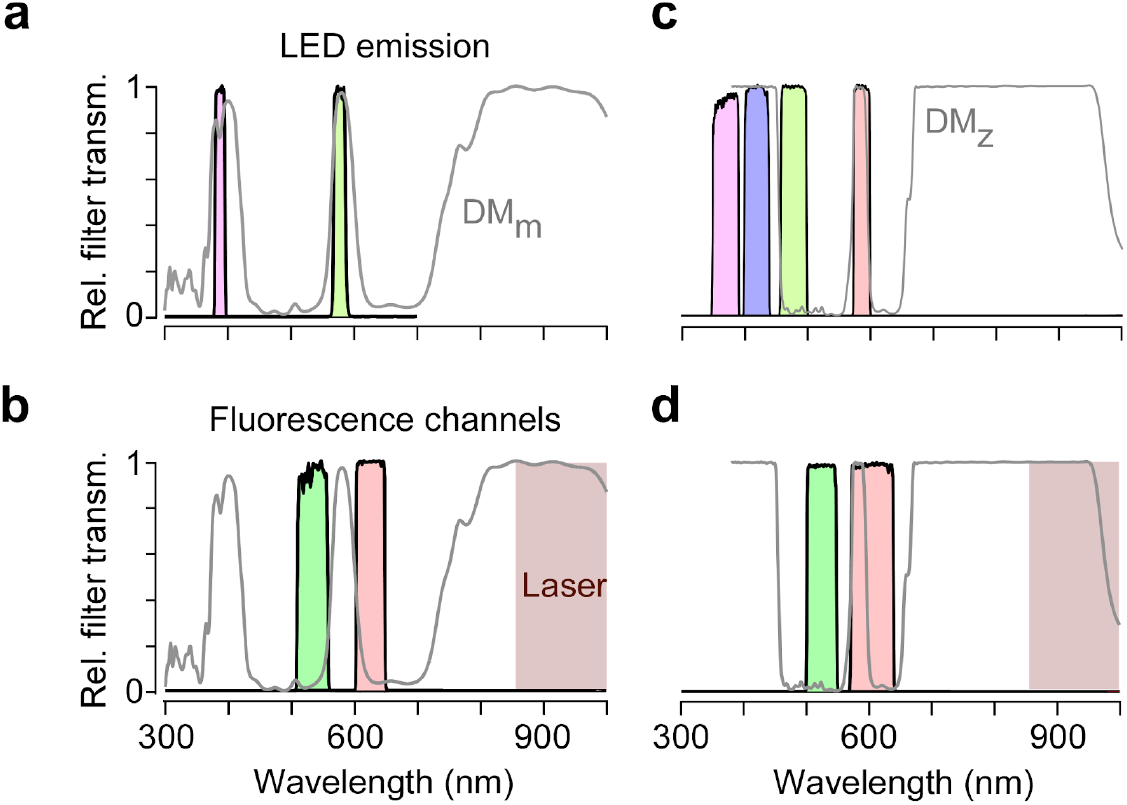
Spectral separation of visual stimulation and fluorescence detection. **a,** Relative transmission of filters in front of UV and green LED as well as of dichroic mirror on top of objective (*cf*. Euler et al., 2009). **b,** Same as (a), for filters in front of PMTs (Methods). Brown shading illustrates the wavelength range used for two-photon laser. **c, d,** Like (a) and (b) but for zebrafish stimulator.

**Supplemental Fig. S3.**
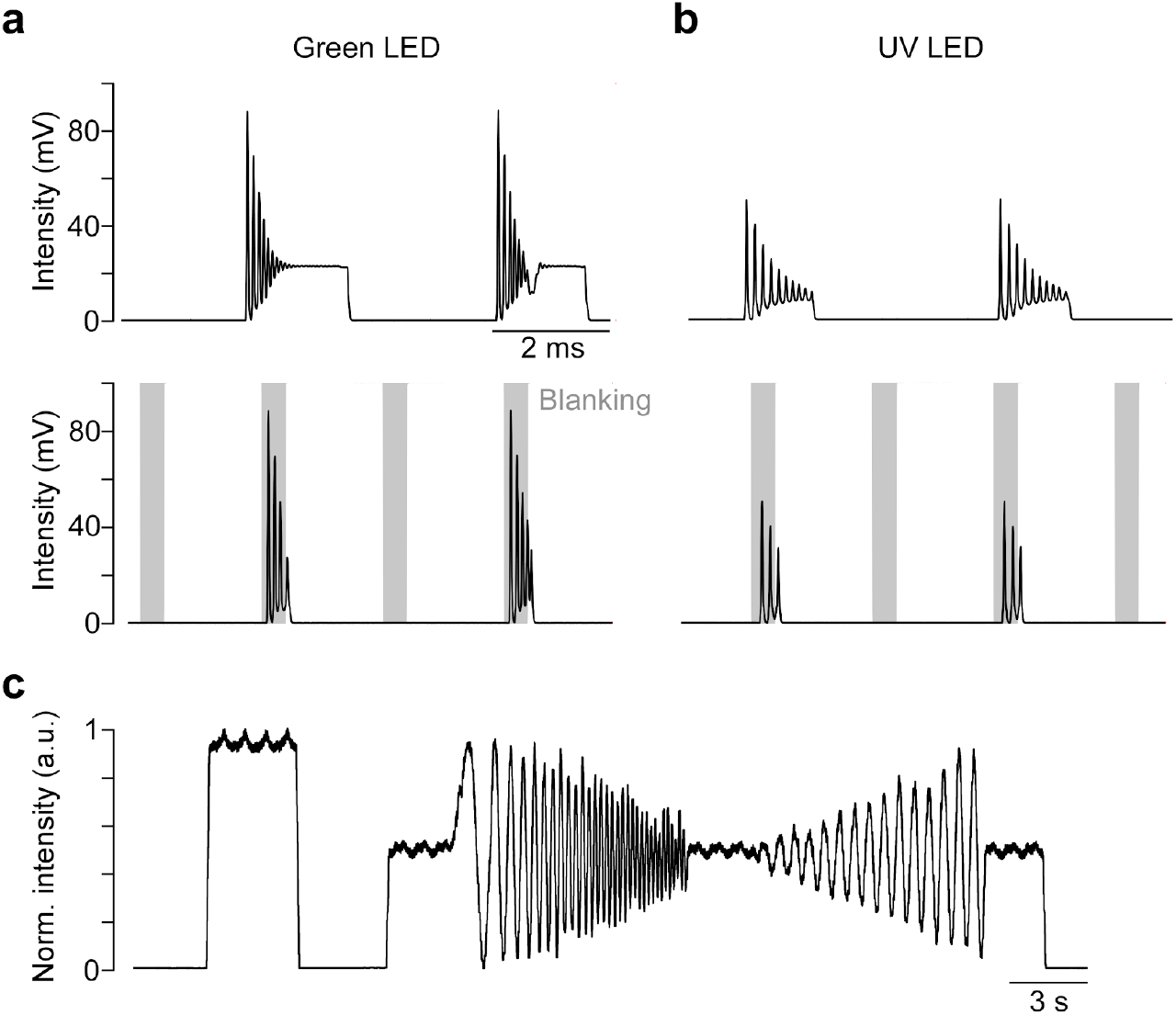
Intensity measurements of the LEDs of the mouse stimulator. **a,** Intensity of green (left) and UV (right) LED measured with a fast photodiode (at 250 kHz; for details, see Methods) without blanking of the LEDs. **b,** Same as (a), but with blanking of the LEDs. Grey shading indicates blanking signal. **c,** Smoothed (box smooth, box width: 100 ms) intensity profile of a full-field chirp stimulus recorded with a fast photodiode.

**Supplemental Fig. S4.**
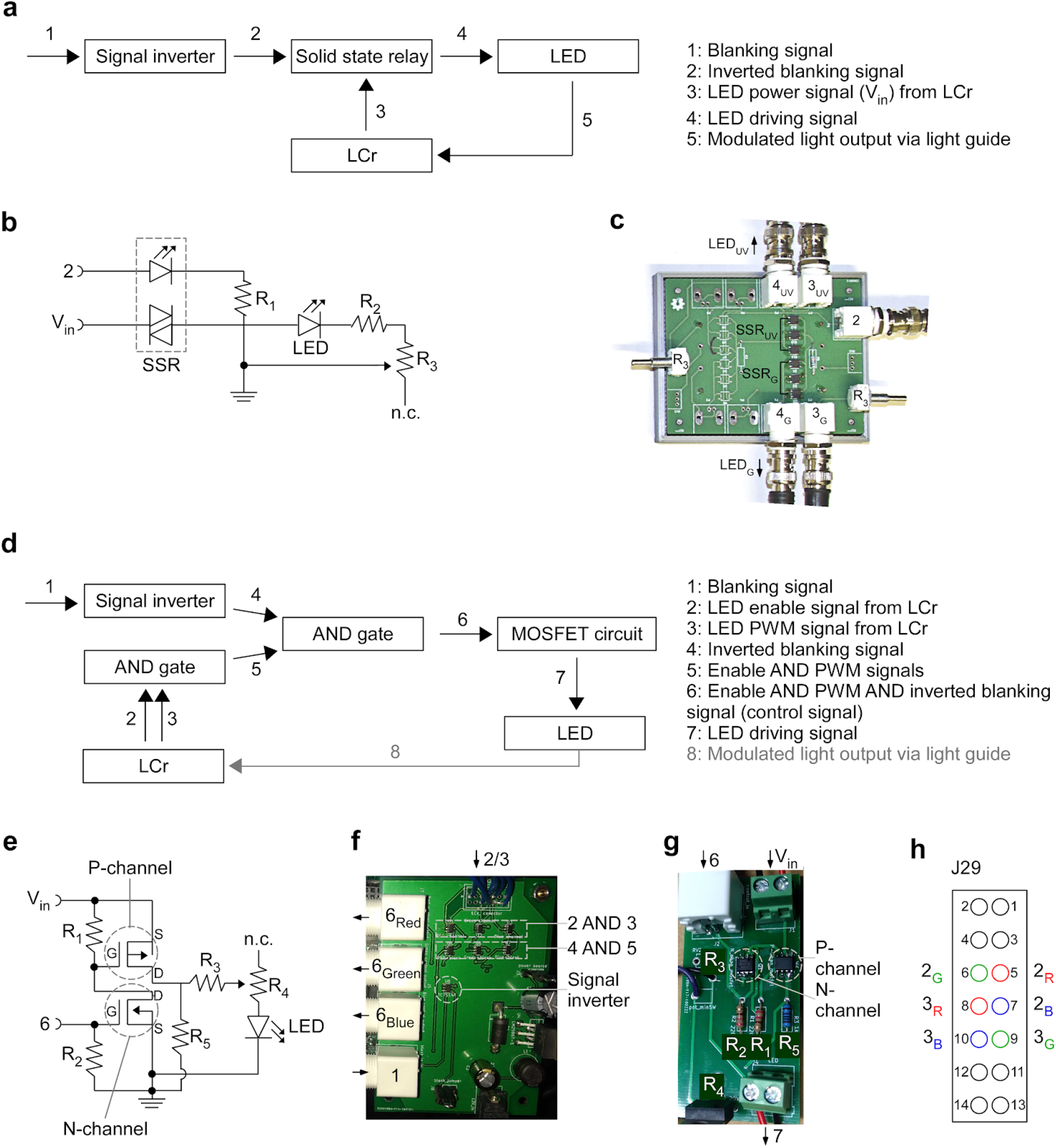
External LED control and temporal separation of stimulation and fluorescence detection. **a,** Schematic illustrating the circuit that ensures that the stimulator LEDs are only on during the microscope′s scanner retrace (for details, see Results). The “laser blanking” signal (1) generated by the scan software is inverted (2) and then used to drive solid-state relays that modulate the LED power signal generated by the LCr (3). This modulated power signal (4) drives the LEDs. The LED light (5) is fed to the LCr via a light guide. **b,** Wiring diagram of the solid-state relay (SSR) circuit (signals like in (a); R_1_=220 Ω, the values for R_2_ and potentiometer R_3_ depend on the LEDs used and typically are in the range of 0-500 Ω). **c,** Picture of the custom-printed circuit board, which can accommodate up to four LED circuits. **d,** Schematic illustrating an alternative circuit to (a-c), where the LEDs are powered from an external supply. Here, the LCr LED control signals (2, 3) go through a logical AND operation and the resulting signal (5) is then combined with the inverted blanking signal (4). The resulting signal (6) is used to switch the LED power using a combination of P and N-channel MOSFETs (*cf.* (e)). Finally, the LED power signal (7) drives the internal or external LEDs. **e,** Wiring diagram of the MOSFET circuit (signals as in (d); R_1_=220 Ω, R_2_=220 Ω, R_3_=0.5 Ω, potentiometer R_4_=25 Ω, R_5_=1 kΩ). **f,g**, Picture of two custom-printed boards responsible for combining the control signals (f) and switching the LEDs (g). **h**, Pinout of connector J29 (“external LED driver connector”) on the LCr board. To disable the LCr’s internal LED drivers, jumper J30 must be installed, while J28 is used to choose between 3.3V or 1.8V supply voltage (see Table 1 for links to LCr instruction manuals).

**Supplemental Fig. S5.**
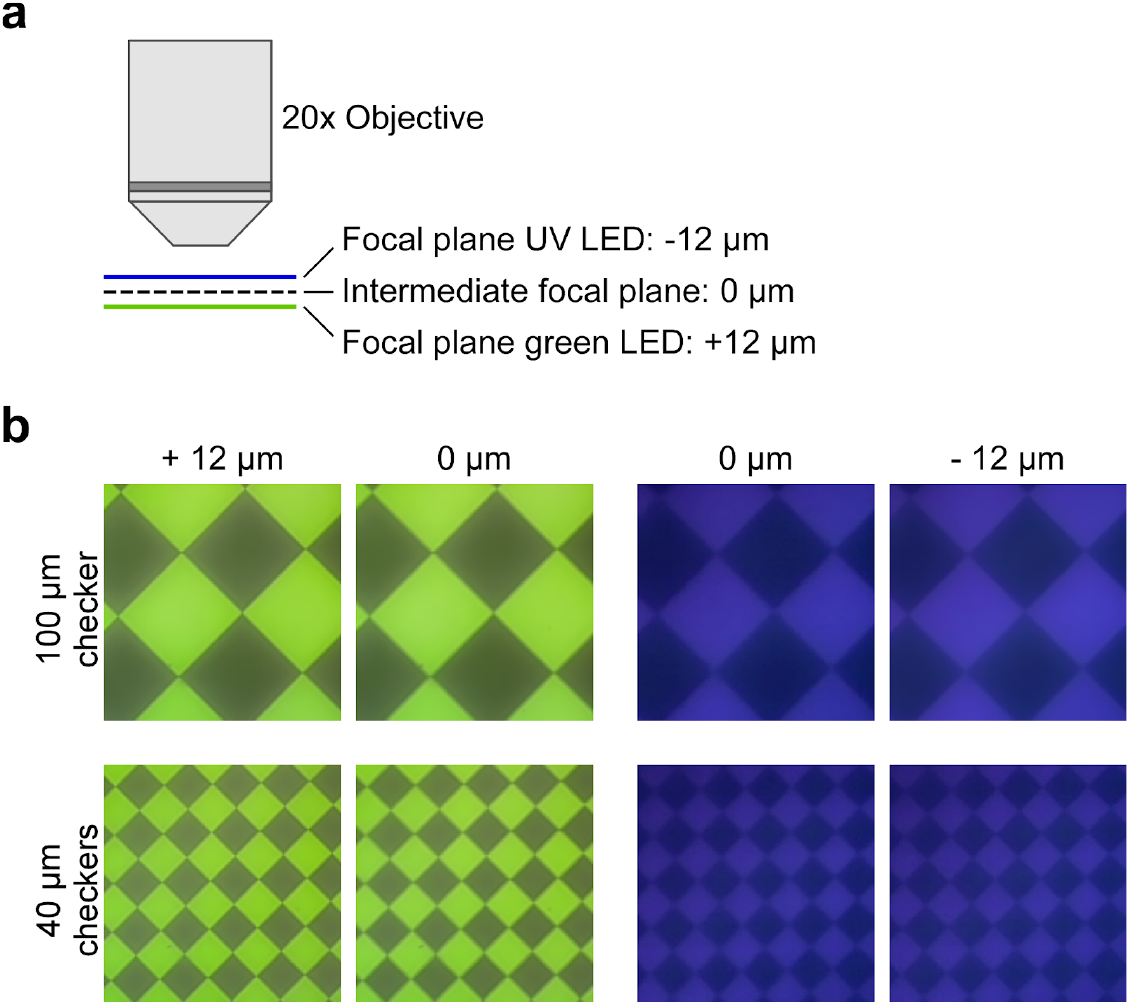
Chromatic aberration of the mouse stimulator. **a,** Schematic illustrating the chromatic aberration-related difference in focal planes of UV and green image in the TTO configuration. **b,** Images of a 100 (top) and 40 (bottom) μm checkerboard stimulus using the green (left) and UV (right) LED, recorded by placing the sensor chip of a raspberry pi camera at the level of the recording chamber. Focus was set to intermediate focal plane (see (a); ±12 μm) or was adjusted for UV and green LED separately (0 μm).

**Supplemental Table S1.** Parts list of the through-the-objective mouse stimulator (cf. Supplemental Fig. S1).

| Part | Description (link) | Company | Item number | Qty. |
| --- | --- | --- | --- | --- |
| LCr | DLP® LightCrafter™ E4500 MKII™ UV(385nm/405nm) + Blue(460nm) + Green(520nm) | EKB Technologies Ltd. | DPM-E4500UVBGMKII | 1 |
| DM 03 | Dichroic mirror (custom made, cf. Supplemental Fig. S1) | AHF Analyse-technik AG | F73-063_z400-580-890 | 1 |
| M 00 | UVFS Plate Beamsplitter | Thorlabs | BSX16 | 1 |
| M 01, 02 | Silver mirrors |  |  | 2 |
| CM | Cold Light Mirror KS 93 / 45° | Qioptiq Photonics GmbH & Co KG | G380255033 | 1 |
| Lens 04 | ACA-254 Achromatic Doublet f = 50 mm | Thorlabs | ACA254-050-A | 1 |
| Lens 05 | AC-508 Achromatic Doublet | Thorlabs | AC508-100-A-ML | 1 |
| BP 03 | Band pass filter 380-407 / 562-589 | AHF Analyse-technik AG | F59-003 | 1 |
| z stage | 13 mm Travel Vertical Translation Stage | Thorlabs | MVS005/M | 2 |
| X or y stage | 13 mm Translation Stage | Thorlabs | MT1B/M | 4 |

